# AcrIF11 is a potent CRISPR-specific ADP-ribosyltransferase encoded by phage and plasmid

**DOI:** 10.1101/2024.08.26.609590

**Authors:** Daphne F. Chen, Leah T. Roe, Yuping Li, Adair L. Borges, Jenny Y. Zhang, Palak Babbar, Sourobh Maji, Maisie G.V. Stevens, Galen J. Correy, Morgan E. Diolaiti, Dominique H. Smith, Alan Ashworth, Robert M. Stroud, Mark J.S. Kelly, Joseph Bondy-Denomy, James S. Fraser

## Abstract

Phage-encoded anti-CRISPR (Acr) proteins inhibit CRISPR-Cas systems to allow phage replication and lysogeny maintenance. Most of the Acrs characterized to date are stable stoichiometric inhibitors. While enzymatic Acrs have been characterized biochemically, little is known about their potency, specificity, and reversibility. Here, we examine AcrIF11, a widespread phage and plasmid-encoded ADP-ribosyltransferase (ART) that inhibits the Type I-F CRISPR-Cas system. We present an NMR structure of an AcrIF11 homolog that reveals chemical shift perturbations consistent with NAD (cofactor) binding. In experiments that model both lytic phage replication and MGE/lysogen stability under high targeting pressure, AcrIF11 is a highly potent CRISPR-Cas inhibitor and more robust to Cas protein level fluctuations than stoichiometric inhibitors. Furthermore, we demonstrate that AcrIF11 is remarkably specific, predominantly ADP-ribosylating Csy1 when expressed in *P. aeruginosa*. Given the reversible nature of ADP-ribosylation, we hypothesized that ADPr eraser enzymes (macrodomains) could remove ADPr from Csy1, a potential limitation of PTM-based CRISPR inhibition. We demonstrate that diverse macrodomains can indeed remove the modification from Csy1 in *P. aeruginosa* lysate. Together, these experiments connect the *in vitro* observations of AcrIF11’s enzymatic activity to its potent and specific effects *in vivo*, clarifying the advantages and drawbacks of enzymatic Acrs in the evolutionary arms race between phages and bacteria.

## Introduction

The evolutionary arms race between bacteriophages and their hosts has resulted in a plethora of bacteria-encoded immune systems and phage-encoded anti-immune factors. Amidst prolific characterization of systems in this arms race, CRISPR-Cas still distinguishes itself from the rest due to its unique adaptive nature. Phages can evade CRISPR-Cas by encoding anti-CRISPR proteins (Acrs), which can fall under one of three generalized mechanisms for inhibition. The first and most common mechanism is characterized by the stable, stoichiometric association of Acr to the CRISPR-Cas complex. Depending on the Acr, this stable binding can inhibit different functions of the complex: target DNA binding, nuclease activity, nuclease recruitment, and so forth^1^. The second mechanism is CRISPR-Cas complex dissociation or degradation^2,3^. The third mechanism is enzymatic, substoichiometric modification of CRISPR-Cas. Several distinct enzymatic modifications have been observed, including crRNA cleavage^4^ ^5^ ^6^, degradation of Type III CRISPR-Cas signaling molecules^7^, and post-translational modifications such as acetylation^8^ ^9^ and ADP-ribosylation^10^.

In contrast to the dozens of stoichiometric Acrs discovered, there are only five biochemically confirmed enzymatic Acrs. They were discovered using methods such as genome fragment screening or guilt-by-association bioinformatics – methods not specifically targeted towards enzymes. For example, AcrIF11 was previously discovered as a neighbor of a gene encoding an anti-CRISPR repressor protein *aca1* and described as a widespread Type I-F anti-CRISPR protein, which enabled the discovery of Cas12 Acrs via guilt-by-association^11^. Although the sequence was too diverged to permit functional assignment upon discovery, a crystal structure revealed similarity to diphtheria toxin, an ADP-ribosyltransferase^10^. Further *in vitro* biochemical assays demonstrated that AcrIF11 modified N250 of Csy1 in the Type I-F Csy complex, which inhibited target DNA binding^10^.

The emergence of enzymatic Acrs, with more likely being annotated using structural conservation aided by AlphaFold and related methods, raises a question: under what conditions is it favorable for a mobile genetic element (MGE) to encode a catalytic Acr as opposed to a stable, stoichiometric Acr? We hypothesize that the substoichiometric activity of an enzymatic Acr would make it more potent at inhibiting CRISPR-Cas function compared to a stable stoichiometric Acr. Furthermore, we hypothesize that enzymatic Acrs must be specific to their target to avoid producing off-target effects that might interfere with the host, especially when encoded by symbiotic prophages or plasmids. Moreover, hyper-potent and specific Acr proteins could be well suited to stabilizing MGE-host symbiosis, to prevent CRISPR-Cas self-targeting^12^.

Here, we find that ADP-ribosyltransferase AcrIF11 surpasses stable stoichiometric Acrs in protecting phage from Csy complex upregulation and multi-spacer phage targeting. We also show that AcrIF11 effectively rescues lysogens from prophage-induced autoimmunity.

Additionally, we establish that AcrIF11 is highly specific to endogenous Csy1/Cas8 in the *P. aeruginosa* intracellular environment using ADP-ribose-specific immunoblotting. The observed potency and specificity likely explain the observed wide distribution across diverse mobile genetic elements. However, as a potential cost to enzymatic mechanisms, we propose that the reversible nature of post-translational modifications would allow for removal by the host. Indeed, we provide proof of principle for this possibility using diverse macrodomain proteins. Our characterization of AcrIF11 illustrates the versatility, specificity, and potency of ADP-ribosylation in the evolutionary arms race between phages and bacteria.

## Results

### The NMR structure of AcrIF11_Pae2_ reveals conserved ADP-ribosylation machinery

Our original motivation for determining the solution structure of AcrIF11 was to leverage the structural information to assign its molecular function. During the time we worked toward that goal, the X-ray structure of a distinct homolog was determined^10^, permitting functional annotation as an ADP-ribosyltransferase and validation of that activity *in vitro*. In addition, the advent of Alphafold and related methods allows for high confidence structure prediction of AcrIF11 homologs. These two advances position us to compare the experimental structures and conduct broader structural analyses based on predictions for the family.

We used NMR spectroscopy to determine a structure of a homolog of AcrIF11, hereafter noted as AcrIF11_Pae2_ to avoid confusion with the homolog used in a previous study^10^ (PDB: 6KYF), that we will call AcrIF11_Pae1._ Despite the low sequence similarity (approximately 27% amino acid sequence similarity) between AcrIF11_Pae1_ and AcrIF11_Pae2_, their beta sheet domains align well, and there are conserved negatively charged residues in positions that have been previously noted as crucial for catalysis ^13^ ^14^ (Fig 1A). Furthermore, upon titration of cofactor NAD, the largest chemical shift perturbations occurred in the beta sheet region, reinforcing the notion that this region, which is also the most structurally conserved region, is functionally important for NAD binding (Fig 1B). Additionally, the negatively charged AcrIF11_Pae2_ residues D146 and E147, which are conserved across representative AcrIF11 homologs (Supplementary Figure 1), experience large and medium chemical shift perturbations (Fig 1B), respectively, highlighting their importance in NAD binding. The NMR structural ensemble of AcrIF11_Pae2_ shows a broader range of RMSD values for the beta sheet region, including the loop where D146 and E147 are located, than the alpha helical region (Fig 1C). This observation of possible loop flexibility is consistent with the need for structural changes to occur upon NAD binding, as observed in previous bacterial ARTs ^15^. An Alphafold3 prediction of AcrIF11_Pae2_ in complex with NAD also predicted that NAD binds in a similar position to what appears in the crystal structure of AcrIF11_Pae1_ (Fig 1D). Although the prediction places the nicotinamide ribose too far away to interact with D146 and E147 (Fig 1D, right inset), the overall configuration of the NAD molecule is similar to what is observed in the crystal structure (Fig 1D, left inset). Further experimental studies would be required to detail the dynamic changes in AcrIF11_Pae2_ upon NAD binding, and how they differ from AcrIF11_Pae1_. The position of AcrIF11_Pae2_ residues in the NAD-binding/catalytic region appear to be conserved compared to AcrIF11_Pae1_ and diphtheria toxin (Fig 1E), although the exact residue identity may not be conserved.

**Figure 1.**
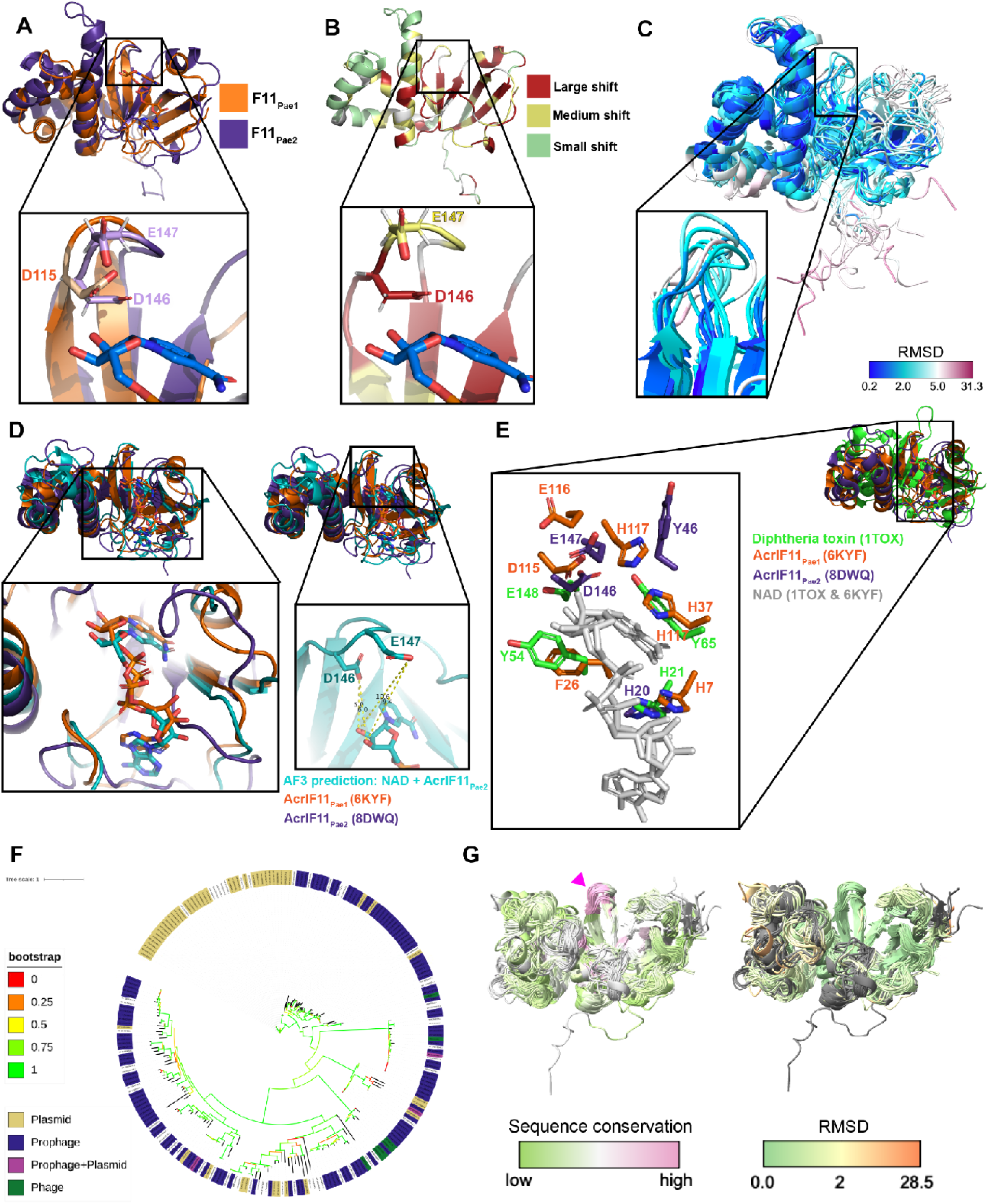
NMR structure of AcrIF11_Pae2_ A) Alignment of AcrIF11_Pae1_ (orange) crystal structure (PDB: 6KYF), and AcrIF11_Pae2_ (purple) NMR structure (PDB: 8DWQ). The inset shows D115 of AcrIF11_Pae1_, a residue previously shown to be important for ART activity^10^, as well as D146 and E147, negatively charged residues of AcrIF11_Pae2_ in a similar position to D115. B) AcrIF11_Pae2_ NMR structure colored according to the magnitude of chemical shift perturbations observed upon NAD (cofactor) titration. The inset shows the same region as the panel A inset. C) AcrIF11_Pae2_ NMR ensemble colored by RMSD. The inset shows the same region as the inset in panel A and B. D) Alignment of AcrIF11_Pae1_ (orange) crystal structure (PDB: 6KYF), AcrIF11_Pae2_ (purple) NMR structure (PDB: 8DWQ), and AcrIF11_Pae2_ (teal) Alphafold3 prediction with NAD. The left inset shows the position of NAD in the AcrIF11_Pae1_ crystal structure in comparison to its position in the Alphafold3 prediction of AcrIF11_Pae2_. The right inset shows positioning of NAD in the Alphafold3 prediction of AcrIF11_Pae2_ relative to the catalytic residues discussed in the panels above. E) Alignment of AcrIF11_Pae1_, AcrIF11_Pae2_, and catalytic domain of monomer of diphtheria toxin (PDB: 1TOX) based on the NAD (light gray) molecules of AcrIF11_Pae1_ and diphtheria toxin. The loop where D146 and E147 are located on AcrIF11_Pae2_ is more extended in the diphtheria toxin, such that there are no diphtheria toxin residues in the equivalent position of Y46. F) Phylogeny of *acrIF11* homologs found after three iterations of PSI-BLAST. Nodes labeled according to the mobile genetic element they are found on: plasmid (gold), prophage (navy), prophage and plasmid (magenta), or phage (green). An enlarged version of this phylogeny can be found in Supplementary Figure 4, and a sample of the alignment used to build the phylogeny is provided in Supplementary Figure 5. G) Alphafold2 predictions of AcrIF11_Pae1_ homologs, colored by sequence conservation and RMSD. Pink arrow indicates the same region as panel A, B, and C insets.

### *acrIF11* is widely dispersed among mobile genetic elements

When *acrIF11* was first discovered, it was difficult to annotate due to its low sequence similarity with functionally annotated proteins. However, structural comparisons with the solved crystal structure permitted identification of ART function^10^. With this annotation in hand, we investigated the extent of *acrIF11* spread (an enlarged version of this phylogeny can be found in Supplementary Figure 4). We found *acrIF11* homologs via PSI-BLAST and queried these representative homologs’ presence on phages and plasmids via a database of bacteriophage and plasmid proteins collated from NCBI. In parallel, we also queried these homologs against a broader provirus and plasmid detector^16^ to account for any unannotated prophage and plasmid regions not present in our homemade database. Combining the results of these two methods, we found that the vast majority of representative *acrIF11* homologs in this phylogeny are present on plasmids and prophages/phages (Fig 1F).

Having observed the structural similarity between AcrIF11_Pae1_ and AcrIF11_Pae2_ despite their low sequence similarity, we were curious if other AcrIF11 homologs displayed a similar structure. Using homologs in the phylogeny in Fig 1F, we clustered homologs with sequence identity cutoff of 95% to prevent redundancy, trimmed the alignment of representative sequences to prevent large gaps in their alignment, and predicted structures of those representative sequences from the alignment using Alphafold2^17^. We used a template date set to earlier than the AcrIF11_Pae1_ deposition date to avoid template bias. The predicted AcrIF11_Pae1_ and AcrIF11_Pae2_ structures aligned well with the experimental structures (Supplementary Figure 2), and most of the structures were predicted with high confidence (Supplementary Figure 3). Alignment of all predicted structures to the predicted AcrIF11_Pae1_ structure revealed that the most variable structural feature is the alpha helical region (approximately residues 51-107 of AcrIF11_Pae1_ PDB 6KYF), while the most structurally conserved region is the loop where AcrIF11_Pae2_ D146/E147 and AcrIF11_Pae1_ D115/E116 are located (pink arrow, Fig 1G). These results show that Alphafold2 is capable of detecting conserved catalytic regions across structures, even when the overall sequence similarity is low.

### ADP-ribosylation from AcrIF11 is specific to Csy1 and required for protecting phage against CRISPR-Cas

Although AcrIF11’s ADP-ribosyltransferase activity has been validated *in vitro*^10^, the specificity of AcrIF11 in the cellular environment is still unknown. To detect the ADP-ribose modification imparted by AcrIF11 on Csy1 (Cas8 homolog), we expressed AcrIF11_Pae2_ on a plasmid in PA14, and then blotted the lysate for ADPr-modified protein with an ADPr-specific antibody (Fig 2A). A previously engineered PA14 strain was used, which has an sfCherry2-fusion to Csy1 at the endogenous location in the genome^18^ to enable us to blot for Cherry for both a loading control and to detect any changes in Csy1 length/stability upon modification. Upon expression of WT AcrIF11_Pae2_, we observed an ADPr band at the expected mass of sfCherry2-Csy1 (Fig 2A, F11_Pae2_WT lane, red arrow). The strong band slightly below 70 kDa is presumably an ADP-ribosylated protein in the PA14 lysate, as shown by its presence in the absence of AcrIF11_Pae2_ (Fig 2A, EV lane). An anti-Cherry immunoblot indeed confirmed equal loading and protein levels of the different samples, confirming that modification does not induce proteolysis. Furthermore, in a strain where CRISPR RNAs will not be expressed and processed, AcrIF11_Pae2_ was unable to ADP-ribosylate Csy1 (Fig 2A, F11_Pae2_WT Δcr1Δcr2 lane), suggesting that the fully assembled Csy complex is necessary for AcrIF11 to recognize and/or bind to Csy1. This observation is in line with previous *in vitro* studies showing that residues on a neighboring protein in the Csy complex, Csy3/Cas7f, are necessary for AcrIF11 activity^10^. Regarding specificity in the intracellular environment, the only detectable covalent modification imparted in an AcrIF11-dependent manner was on Csy1, suggesting that this is a very specific anti-CRISPR mechanism.

**Figure 2.**
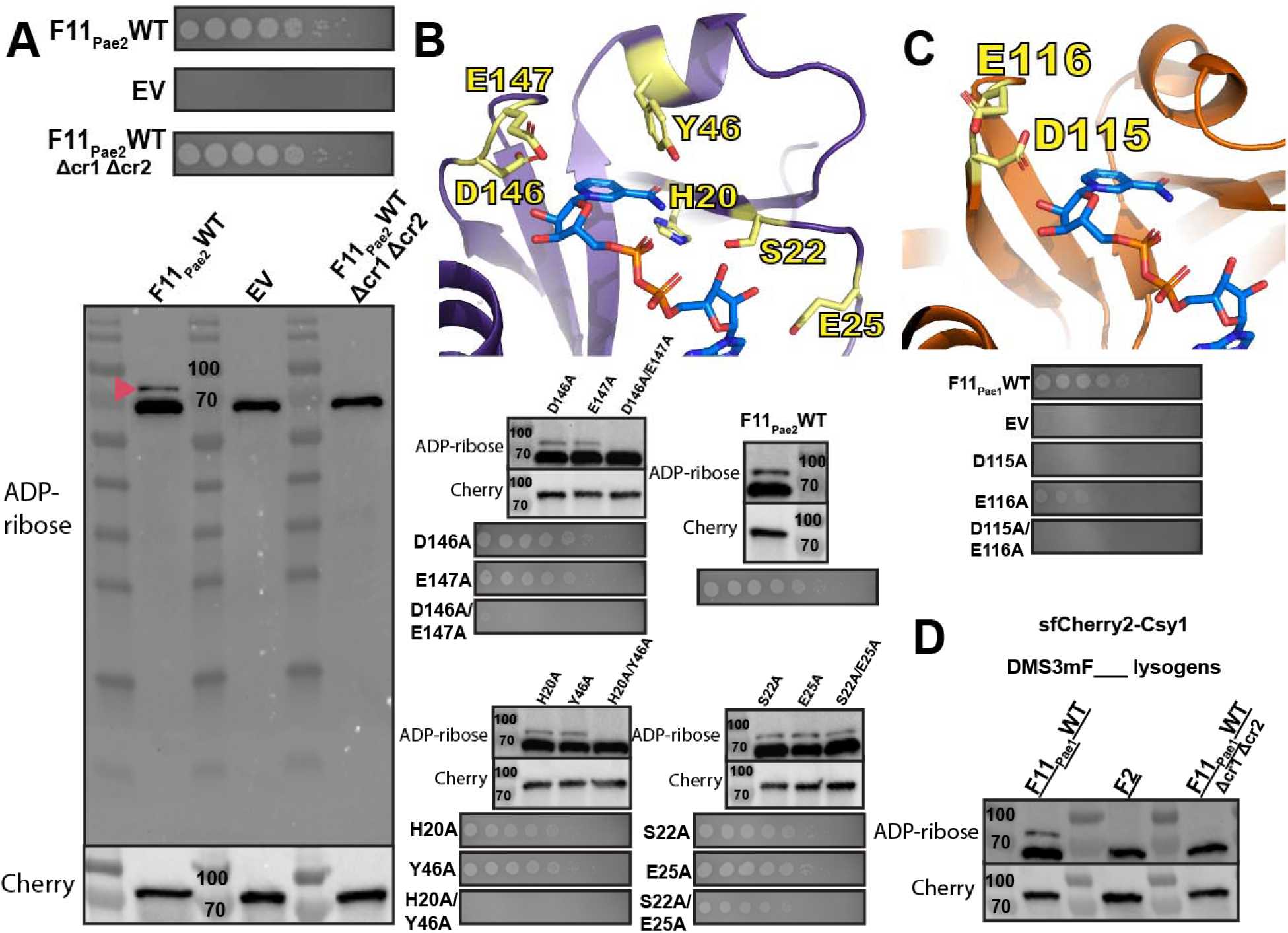
ADP-ribosylation activity of AcrIF11 A) Plaque assays of PA14 chromosomally expressing sfCherry2-Csy1 and also containing a plasmid expressing wildtype AcrIF11_Pae2_ (F11_Pae2_WT) or empty vector (EV). Δcr1Δcr2 indicates PA14 with both CRISPR arrays deleted. These strains were also lysed and probed for ADPr and Cherry. Pink arrow indicates sfCherry2-Csy1 protein modified with ADPr. B) Plaque assays and Western blots as in panel A, but with wildtype and mutant versions of AcrIF11_Pae2_. Mutants are colored yellow on the AcrIF11_Pae2_ NMR structure (PDB: 8DWQ). The NAD molecule is from alignment with the AcrIF11_Pae1_ crystal structure (PDB: 6KYF). C) Plaque assays and Western blots as in panel A, but with wildtype and mutant versions of AcrIF11_Pae1_. Mutants are colored yellow on the AcrIF11_Pae1_ crystal structure (PDB: 6KYF). D) Western blots of lysate from PA14 sfCherry2-Csy1 DMS3m lysogens encoding the indicated Acr. AcrIF2 is a stable stoichiometric binder with no enzymatic activity.

In order to directly correlate ADP-ribosylation to successful CRISPR-Cas inhibition and phage replication, we tested AcrIF11_Pae2_ mutants’ ability to protect CRISPR-targeted DMS3m, and also detected ADP ribosylation in the lysate (Fig 2B). The D146/E147 pair was chosen for its conservation across homologs as well as the potential importance of their negative charge in stabilizing the positively charged transition state of NAD^13^ ^14^. The H20/Y46 pair was chosen because the structure suggests they are important residues for stabilizing the nicotinamide leaving group. Lastly, we hypothesized that the S22/E25 pair were important for NAD binding.

For D146A/E147A and H20A/Y46A, disappearance of ADPr in lysate is matched by the absence of phage replication, showing that without AcrIF11’s ADP-ribosylation of Csy1, CRISPR-Cas activity has targeted the phage. We also mutated the equivalent catalytic residues based on the AcrIF11_Pae1_ crystal structure 6KYF (Fig 2C) and observed no CRISPR inhibition in D115A, consistent with the absence of ADPr from *in vitro* Western blot in a previous study^10^.

Lastly, the S22A/E25A mutant in AcrIF11_Pae2_ still inhibited CRISPR-Cas function and appended an ADPr-modification, demonstrating that the residues are redundant for Acr catalytic activity.

Having observed successful ADP-ribosylation of Csy1 using plasmid-expressed AcrIF11_Pae2_, we were curious if native protein expression levels would also result in robust ADP-ribosylation. To investigate this possibility, we made a PA14 sfCherry2-Csy1 DMS3mAcrIF11_Pae1_ lysogen.

We first engineered the DMS3m phage to express *acrIF11* from the native anti-CRISPR locus, replacing the endogenous *acrIE3* gene via recombination. This approach was similarly used previously to compare distinct Acr proteins in an otherwise isogenic phage background^19^. Using the PA14 sfCherry2-Csy1 strain with a DMS3mAcrIF11_Pae1_ prophage, we observed an ADP-ribose band at the mass of the sfCherry2-Csy1 fusion (Fig 2D, lane F11_Pae1_WT). As a negative control, we used a previously constructed DMS3m phage with AcrIF2 and made a lysogen; AcrIF2 is a stable stoichiometric Acr that targets the same region of the Csy complex but has no ADP-ribosyltransferase activity. As expected, we saw no ADP-ribose band at the sfCherry2-Csy1 mass (Fig 2D, lane F2). To test if assembly of the Csy complex is important as it was previously (Fig 2A F11_Pae2_WT Δcr1Δcr2 lane), we made a PA14 Δcr1Δcr2 sfCherry2-Csy1 DMS3mAcrIF11_Pae1_ lysogen. In line with our previous observations, we did not see an ADP-ribose band at sfCherry2-Csy1 mass. These results show that even under a native promoter where protein expression is more than an order of magnitude less than plasmid expression^20^, AcrIF11’s substoichiometric nature allows it to ADP-ribosylate and inactivate the Csy complex.

### The substoichiometric activity of AcrIF11 protects phages in situations where stoichiometric Acr activity fails

In addition to our observation of *acrIF11* homologs in diverse MGEs (Fig 1E), we also observed that *acrIF11* is far more widespread than other type I-F Acrs (Fig 3A). *acrIF11* is encoded by MGEs present in over 35 bacterial genera, a full list of which can be found in Supplementary Table 2. These findings suggest that *acrIF11* is a versatile Acr that confers a fitness advantage in many environments and/or niches. We hypothesized that post-translational modifications are a potent mechanism of rapid and complete CRISPR-Cas inhibition, perhaps explaining the widespread nature of *acrIF11*. Borges et. al. have previously shown that Acr proteins vary in potency during phage infection. For “strong” Acr proteins such as AcrIF1, a low concentration is needed for both successful lytic infection and establishment of lysogeny. In contrast to AcrIF1, “weak” Acrs such as AcrIF4 must be in relatively higher concentration for successful infection^19^. When expressed from a phage infecting PA14, AcrIF11_Pae1_ allowed for robust phage replication across a wide range of input concentrations similar to that of AcrIF1, inducing culture lysis at

**Figure 3.**
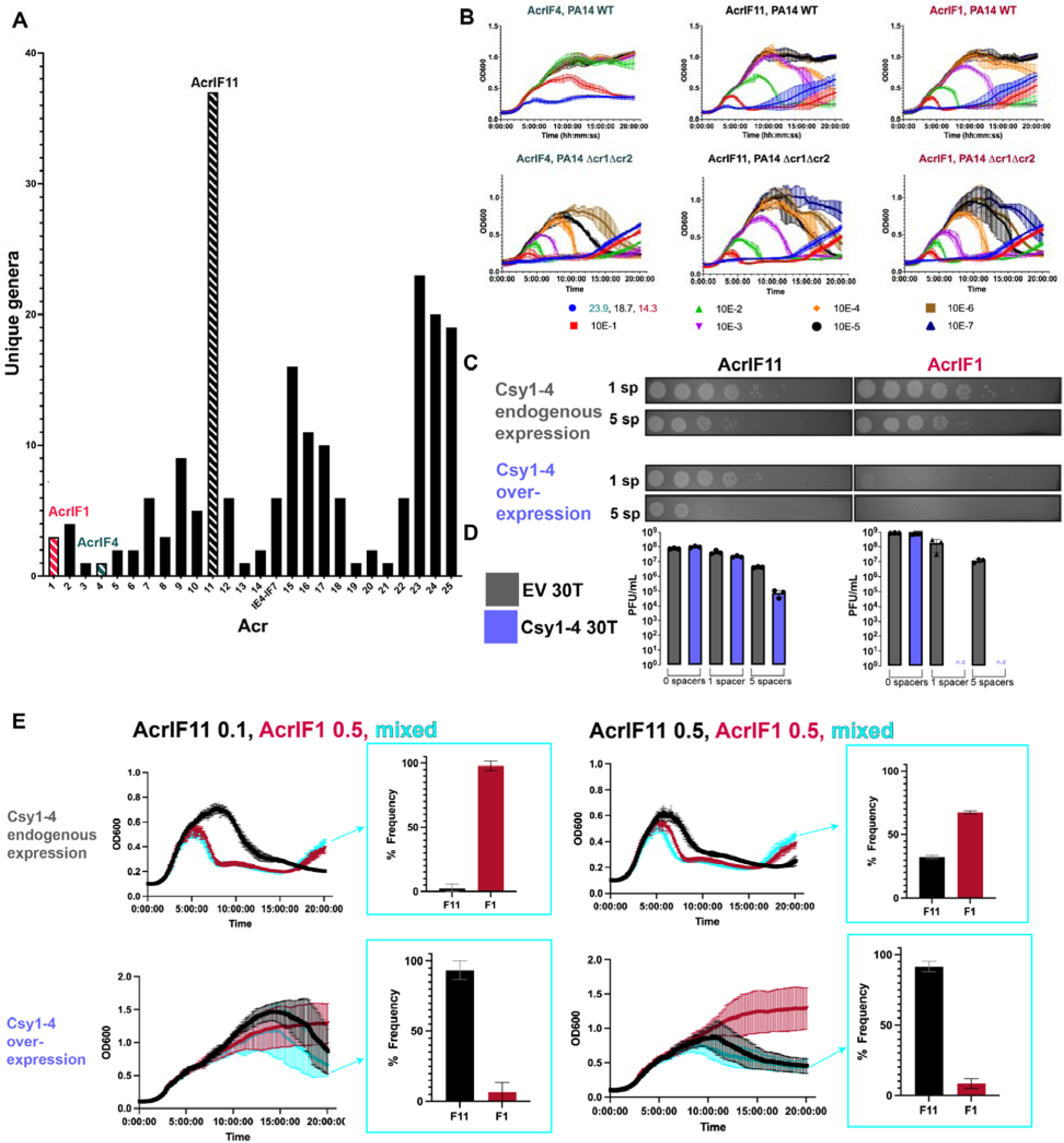
*acrIF11* distribution and protection of lytic phage. A) Plot of the number of bacterial genera that each Type I-F Acr can be found in, via BLASTp. Acrs of interest have been marked with striped bars: AcrIF11 (black), AcrIF1 (pink), AcrIF4 (teal). B) Liquid growth curves of PA14 WT (1 spacer targeting DMS3mvir) infected by DMS3mvir encoding the indicated Acr, across a range of multiplicities of infection (MOI). Plots are the average of three biological replicates. The second row of plots labeled “Δcr1Δcr2” are liquid growth curves of PA14 Δcr1Δcr2 (no spacers targeting DMS3mvir) infected by the same phage as above. C) Plaque assays of PA14 spotted with 10-fold serial dilutions of DMS3m phage encoding either AcrIF11_Pae1_ (left column) or AcrIF1 (right column). PA14 strains encoded either 1 spacer (wildtype targeting) or 5 spacers (high targeting) in their CRISPR arrays targeting DMS3m; each spacer condition was also combined with endogenous chromosomal Csy1-4 expression or overexpressed Csy1-4 on pHERD30T induced with 0.1% arabinose. Plaque assays are representative of three biological replicates. D) Quantification of full plate infections using the same strains and phage as panel C. E) Phage competition growth curves, using the same wildtype and high targeting backgrounds of PA14 as in panel C. PA14 strains were infected with DMS3mvir phages encoding the indicated Acr. Cyan curves show PA14 strains infected with both DMS3mF11vir and DMS3mF1vir at the indicated MOI. The resulting frequencies of observing AcrIF1 or AcrIF11 by genotyping at the end of the time course are plotted. Growth curves and frequencies are the average of three biological replicates.

MOIs as low as 1.87E-5 (Fig 3B). This shows that AcrIF11_Pae1_ exhibits strong inhibition of type I-F CRISPR-Cas during lytic infection, although this experiment does not distinguish between stoichiometric vs substoichiometric activity. Interestingly, during infection with exceedingly low MOIs, the strain lacking CRISPR arrays (Δcr1Δcr2) succumbed to lysis at phage concentrations that failed to lyse when CRISPR was intact. Therefore, it remains clear that successful inhibition of CRISPR-Cas activity, whether by an enzyme or a stoichiometric binder, still requires a critical threshold of anti-CRISPR protein, which is determined by the phage population size^19^ ^21^.

Having established that AcrIF11 and AcrIF1 are both strong Acrs, we hypothesized that AcrIF11’s substoichiometric enzymatic activity might allow it to overcome “high pressure” scenarios such as during CRISPR-Cas upregulation or when multiple spacers target the same phage. Recent work has revealed mechanisms for CRISPR-Cas systems to increase Cas protein expression levels after ‘detecting’ an Acr protein ^22^ ^23^. To test this idea, we overexpressed Csy1-4 in PA14 backgrounds with 1 (wildtype) or 5 (high) CRISPR spacers targeting the phage carrying the Acr. AcrIF11_Pae1_ and AcrIF1 allowed phage to replicate at similar levels in both targeting conditions under endogenous levels of Csy1-4 expression.

However, with additional Csy1-4 overexpression, phages encoding AcrIF1 were completely unable to replicate in either the 1 or 5 spacer targeting condition, while phages encoding AcrIF11_Pae1_ were still able to replicate in both the 1 and 5 spacer targeting conditions (Fig 3C, 3D). This suggests that despite both Acr proteins binding to the Csy complex to exert their inhibition, the transient binding and catalytic activity of AcrIF11 allows phage to overcome increases in target concentration through a substoichiometric mechanism.

Next, to directly examine the fitness advantage that AcrIF11 might confer, we conducted a phage competition experiment where phages encoding either AcrIF11_Pae1_ or AcrIF1 were mixed together in equal abundance (Fig 3E, right half) or with AcrIF1 in 5-fold excess of AcrIF11_Pae1_ (Fig 3E, left half). Under wildtype targeting (1sp) and endogenous Csy expression conditions, AcrIF1 was the most frequently occurring Acr in the population at the end of the competition period (Fig 3E, upper half), in both the equal abundance and 5-fold excess conditions. However, under Csy overexpression conditions, AcrIF11_Pae1_ became the most frequently occurring Acr in the population (Fig 3E, lower half) after inputs of both equal abundance and 5-fold AcrIF1 excess conditions. Remarkably, even when AcrIF1 is in 5-fold excess of AcrIF11_Pae1_, it is AcrIF11_Pae1_ that becomes the dominant Acr in the phage population under Csy overexpression (Fig 3E, lower left quadrant). Furthermore, the difference between AcrIF1 and AcrIF11_Pae1_ frequency under Csy overexpression is much larger than the difference between AcrIF1 and AcrIF11_Pae1_ frequency under Csy endogenous expression (Fig 3E lower right quadrant vs Fig 3E upper right quadrant). This indicates that under endogenous levels of Csy expression, phages can use either AcrIF11_Pae1_ or AcrIF1 to inhibit the Csy complex, but when Csy expression levels are increased, it is only AcrIF11_Pae1_ that is robust enough to inhibit Csy and allow for phage replication. Overall, our experiments show that environmental pressures such as increased Cas protein overexpression and multi-spacer targeting can select for the substoichiometric mechanism of AcrIF11_Pae1_ over the more conventional stable binding mechanism of AcrIF1.

### AcrIF11 is a strong Acr that prevents self-targeting

In addition to protecting phage during lytic infection, Acr proteins can be vital for stabilization of the lysogenic state. Rollie et. al. observed autoimmunity between PA14 type I-F CRISPR-Cas and DMS3 prophages that manifested as a growth defect, demonstrating the necessity of factors that mediate phage-host symbiosis by alleviating autoimmunity^12^. In addition to spacers present in the CRISPR array *a priori*, priming spacer acquisition can also lead to new spacers that would perfectly target a self prophage. To test AcrIF11’s ability to prevent prophage-targeted autoimmunity, we conducted growth experiments with PA14 DMS3m lysogens encoding Acrs. The DMS3m phage is targeted by PA14’s endogenous CRISPR array, such that integration into the genome during lysogeny results in self-targeting by CRISPR-Cas. DMS3m phage expressing AcrIF4 was able to establish lysogeny, but a growth defect emerged demonstrating self-targeting over time. Lysogens expressing AcrIF1 and AcrIF11_Pae1_, however, displayed growth comparable to a Δcr1Δcr2 control (Fig 4A), demonstrating full protection. To address the potential for priming spacer acquisition over time, we passaged the lysogens for three days but observed no growth defect in the presence of AcrIF11 and AcrIF1. These results show that both AcrIF11 and AcrIF1 are strong inhibitors of CRISPR-Cas in the lysogenic cycle, stabilizing a self-targeted MGE.

**Figure 4.**
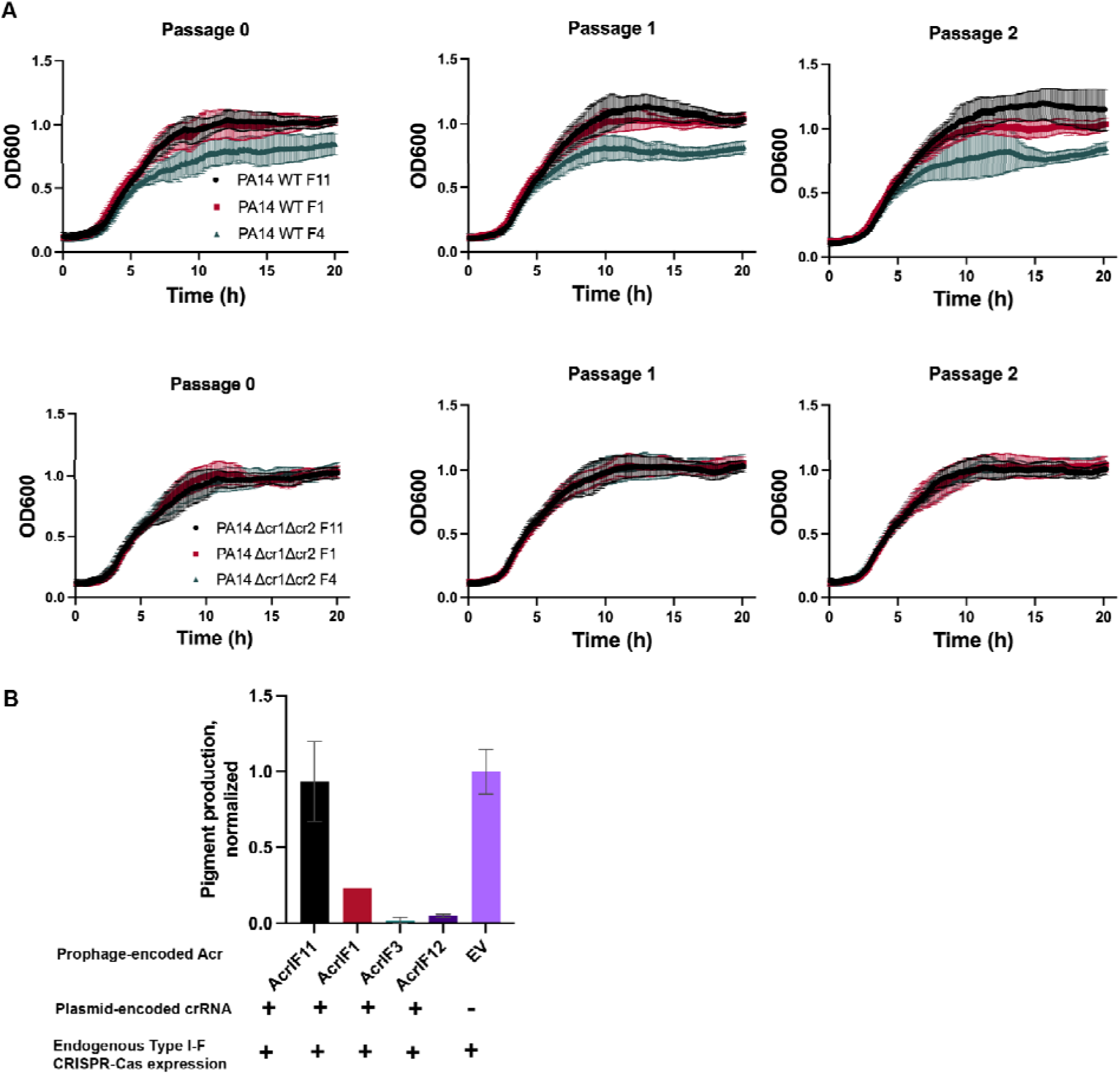
Prophage-encoded *acrIF11* prevents self-targeting A) Liquid growth curves of PA14 lysogens with a DMS3m prophage encoding either AcrIF11_Pae1_, AcrIF1, or AcrIF4, as indicated in the legend. “WT” refers to wild type PA14, and “Δcr1Δcr2” refers to PA14 with both CRISPR arrays deleted. Plots are the average of three biological replicates. B) CRISPRi experiment assessing self-targeting via pyocyanin production. Pyocyanin production was measured after growth of PA14 DMS3m lysogens containing a plasmid encoding *phzM*-targeted (pyocyanin synthesis gene-targeted) crRNA, with DMS3m encoding the Acr indicated on the x axis.

To quantitatively query the *in vivo* self DNA binding ability of the Csy complex as a proxy for self-targeting risk, we assessed the degree of CRISPR interference (CRISPRi) enacted by a crRNA targeting *phzM*, a gene involved in pyocyanin synthesis. Repression of *phzM* silences pyocyanin production, a colored pigment, while de-repression by lysogen-encoded Acrs restores pyocyanin production (Fig 4B). While AcrIF1 has been previously shown to disable CRISPRi, overexpression of the *phzM-*targeting crRNA overwhelmed this Acr, leading to repression of pigment production. In a previous study, AcrIF4 was also shown to partially inhibit CRISPR-Cas^24^ in the same CRISPRi setup, as AcrIF1 did in our experiment. Unlike AcrIF1 and AcrIF4, AcrIF11 fully disabled the Csy complex under overexpression of the *phzM-*targeting crRNA, allowing full pyocyanin production compared to a strain with an non-targeting crRNA. This shows that, while AcrIF11 and AcrIF1 are capable of stabilizing lysogeny in wildtype targeting conditions (Fig 4A), only AcrIF11 can prevent Csy from stably binding to its own genome under “high” targeting conditions. Thus, AcrIF11 activity is fully capable of inhibiting Csy complex from stably binding to its own genome, likely an important aspect of the biology of this MGE-encoded Acr.

### ADP-ribosylation of Csy1 is reversible

Lastly, we hypothesized that enzymatic modifications such as ADPr could be removed by enzymes known as macrodomains, or host-encoded eraser enzymes (Fig 5A) as a continuation of the phage-bacteria arms race. Macrodomains have previously been found in defense and anti-defense contexts; for example, DarG of the DarTG toxin/anti-toxin system can remove ADPr from ADP-ribosylated DNA^25,26^, and ThsA of the Thoeris system can cleave NAD into nicotinamide and ADPr^27^. However, neither DarG nor ThsA have been shown to remove ADPr from a protein and little is known about *Pseudomonas* macrodomain proteins. To investigate the possibility of ADPr removal from Csy1 via macrodomain, we introduced purified macrodomains into lysate from our sfCherry2-Csy1 strain expressing plasmid-encoded AcrIF11_Pae2_. Due to ongoing interest in one of our labs (J.S.F.) in macrodomain activity in the context of human viral infections, the macrodomains we chose to introduce were from Eastern Equine Encephalitis Virus (EEEV), Barmah Forest Virus (BFV), and humans (hMacroD2). Compared to our control lysate that did not receive any macrodomain (Fig 5B, solid red arrow), we observed disappearance of the ADPr signal from sfCherry2-Csy1 (Fig 5B, hollow red arrow), and from the background ADP-ribosylated protein (∼70 kD), with hMacroD2 added in. The ADPr signals from incubation with BFV and EEEV appeared to be the same as our control, indicating no removal of ADPr.

**Figure 5.**
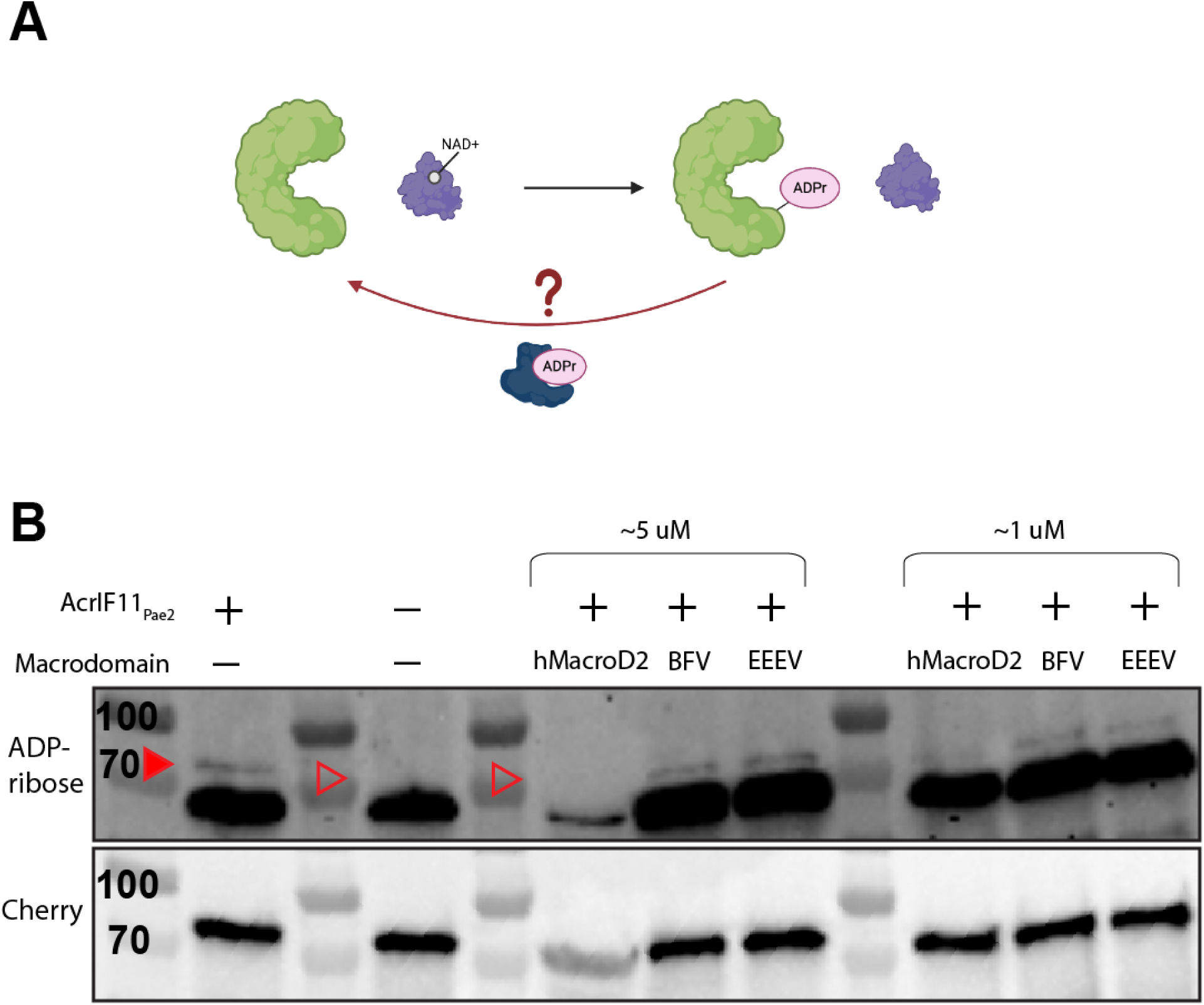
Macrodomain removal of ADPr. A) Schematic of Csy complex (green) modified by AcrIF11 (purple) using NAD as a cofactor. We hypothesize that a macrodomain (dark blue) could remove ADPr from the Csy complex. B) Western blot of purified macrodomain incubated with lysate from PA14 sfCherry2-Csy1 overexpressing AcrIF11_Pae2_ on a plasmid. The mixture was blotted for ADP-ribose and Cherry as in Figure 2. Quantification of the band intensities is available in Supplementary Figure 7. The sfCherry2-Csy1 ADPr band is indicated by the solid red arrow, and its absence is indicated by the hollow red arrows. We believe the decrease in Cherry intensity in the hMacroD2 5 uM lane is not due to decreased loading, but rather the similar molecular weight of our hMacroD2 construct and sfCherry2-Csy1 leading to overcrowding in that molecular weight range as observed by Ponceau stain in Supplementary Figure 8, and thus decreased antibody binding.

To test the activity of these macrodomains, we used a phage encoding AcrIF11 to infect PA14 overexpressing the macrodomains in a liquid infection setup. However, we were unable to observe increased bacterial growth that would indicate antagonism of the anti-CRISPR mechanism compared to our negative control (Supplementary Figure 9). This could be due to potential low/no expression of these evolutionarily distant macrodomains in the cellular environment of PA14. Despite this, our observation of purified macrodomain removing ADPr in lysate shows that the ADPr on Csy1 is accessible and able to be removed *in vitro*. Macrodomain specificity, activity levels, and native substrates are an active field of exploration, so it is difficult to reach a conclusion about why hMacroD2 was able to remove ADPr while BFV and EEEV were not. Although these results are not definitive, they open the possibility of another aspect of the phage-bacteria arms race: host-encoded erasers of enzymatic Acr modifications. We hypothesize that the single-residue specificity of AcrIF11’s modification and its transient binding to the Csy complex could be a disadvantage in strains that encode a modification eraser, compared to a stoichiometric binder such as AcrIF1, which is stably bound to a larger interface of the Csy complex.

## Discussion

In this study we investigated the cellular behavior of the ADP-ribosyltransferase AcrIF11 and found that it confers several benefits to a phage’s ability to infect its host. We showed that enzymatic Acrs outperform stoichiometric Acrs when the phage faces a “high pressure” scenario, such as increased targeting from multiple spacers and Csy upregulation. Additionally, we observed AcrIF11’s potency in lysogeny by measuring the extent to which it prevents host Csy from self-targeting a prophage in the host genome. We also demonstrated that AcrIF11 can rescue lysogens from growth defects due to autoimmunity resulting from prophage targeting.

Furthermore, we established that AcrIF11 is highly specific in its host’s intracellular environment – presumably so it does not interfere with activity from host enzymes (Figure 6). AcrIF11’s potency and specificity illustrates its versatility as an Acr, which is corroborated by our phylogenetic analysis showing it is the most widespread Type I-F Acr and can be found in many distinct mobile genetic elements.

**Figure 6.**
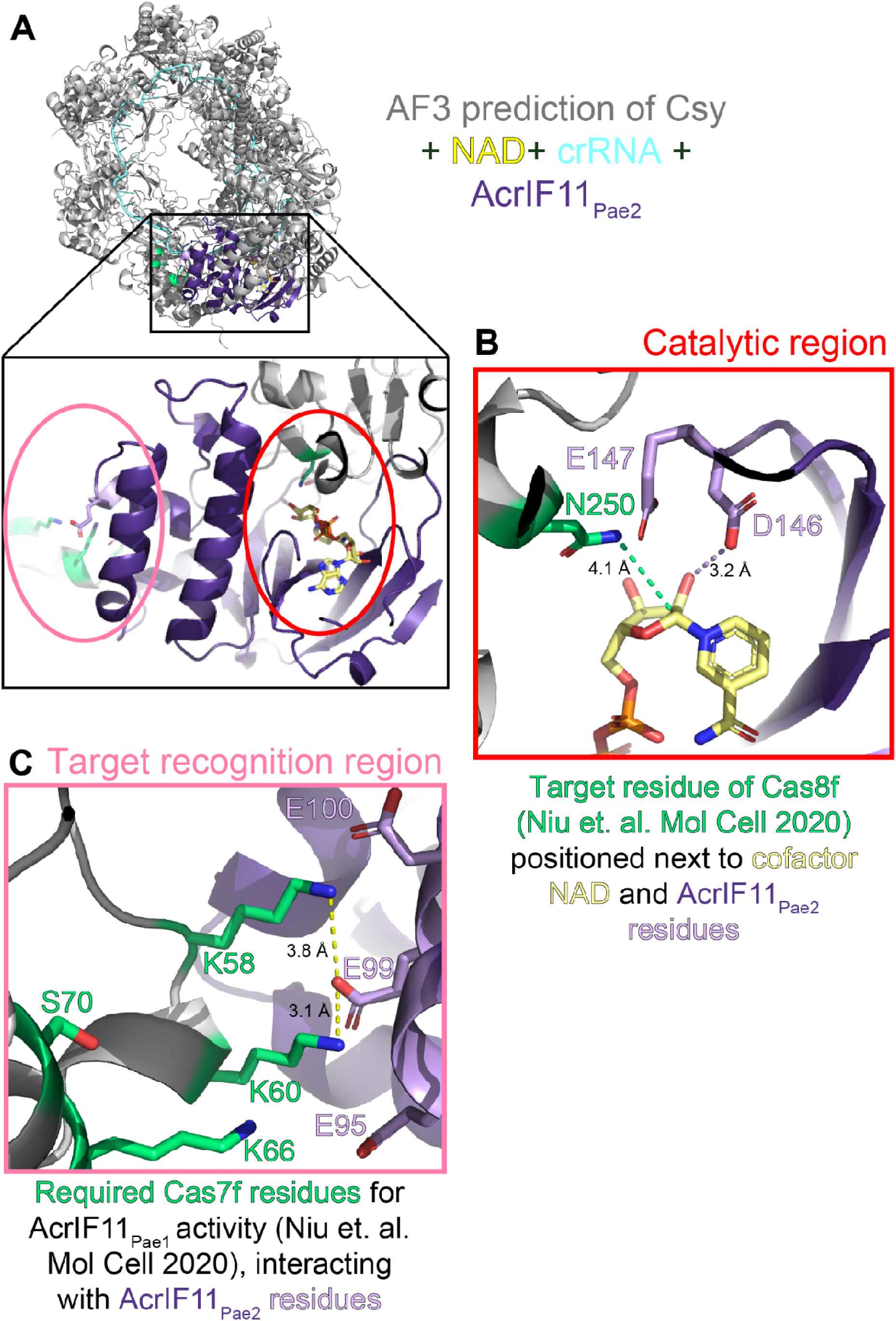
Alphafold3 prediction of AcrIF11-Csy recognition interface A) Alphafold3 prediction of Csy complex (gray) with crRNA (blue, sequence from PDB 6B45), NAD (yellow), and AcrIF11_Pae2_ (purple). Scores of iPTM = 0.69 and pTM = 0.72 indicate this is a medium confidence prediction.^28^ ^29^ The inset depicts the overall position of AcrIF11_Pae2_ on the Csy complex, as well as the main two interfaces of interest: the catalytic region where NAD is bound, and the target recognition region. B) Catalytic region of AcrIF11_Pae2_ with NAD-interacting residues D146 and E147 highlighted in light purple. The dashed purple line indicates a potential hydrogen bond in this static structure, although the dynamics of catalysis could place D146 and E147 closer. N250, the target residue identified by a previous study^10^, is highlighted in green. The green dashed line is the distance between atoms that would be covalently linked after the ADP-ribosylation reaction. C) Target recognition region between AcrIF11_Pae2_ and the Csy complex. Csy3/Cas7f residues that have been previously identified as important for AcrIF11_Pae1_ activity are highlighted in green. The dashed yellow lines indicate potential charged interactions between Csy3/Cas7f and AcrIF11_Pae2_.

AcrIF11’s enzymatic activity as an ADP-ribosyltransferase was discovered previously via crystal structure determination and further validated with *in vitro* biochemistry. Had structure prediction been as widely available as it is now, AcrIF11’s enzymatic activity could have been hypothesized after running a predicted structure through DALI and noticing structural similarity to diphtheria toxin. Our Alphafold predictions of AcrIF11 homologs showed that catalytic sites could be predicted, presumably because catalytic sites are structurally conserved across many enzymes in the PDB. We envision structure prediction and comparison to play a larger role in determining enzymatic Acr function in the future; the functional hypotheses that structure prediction provides could narrow the search space for a protein of unknown function, leading to faster experimental validation and iteration.

We found that numerous mobile genetic elements encode AcrIF11 homologs. Our structure of one particular homolog AcrIF11_Pae2_, in conjunction with Alphafold2 predicted structures of the homologs from our phylogeny, demonstrates that dissimilar amino acid sequences from across the sequence space of AcrIF11 homologs can still converge on a common catalytic ART fold.

That being said, the details of binding within the ART fold are currently difficult to classify for the AcrIF11 family due to small sample size. Although AcrIF11_Pae1_ appears to be most structurally similar to diphtheria-like ARTs, its key residues appear to be H-F-H-D^10^ instead of the canonical H-Y-Y-E^30^. Furthermore, the exact classification of AcrIF11_Pae2_ cannot be determined from our NMR structure alone because this structure was not solved in complex with NAD. The details of NAD-AcrIF11_Pae2_ binding remain unknown, but our mutagenesis of AcrIF11_Pae2_ suggests it is similar to diphtheria-like ARTs. The chemical shifts of AcrIF11_Pae2_ measured upon NAD titration illustrate the importance of loops analogous to the A- and D-loops near the catalytic site, in line with previous observations of these loops’ importance^31^.

An unanswered question about AcrIF11, and the ART family more broadly, is how target recognition occurs. Because the catalytic loops are known to be important for both catalysis and target recognition^31^, more *in vitro* studies will be needed to determine if catalysis and Csy1 recognition residues can be disentangled for AcrIF11, or if the exact same set of residues plays both roles. Mutagenesis has shown that residues in Csy3/Cas7f are required for AcrIF11_Pae1_ ART activity on N250 of Csy1^10^; whether these Csy3 residues interact with an AcrIF11 loop residue in the beta sheet region, or a residue further away in the alpha helical region, remains to be seen. We observed the alpha helical region of AcrIF11 to be mostly poorly conserved and experience little to no chemical shift perturbations upon NAD titration. However, whether that is because this region solely plays a role in Csy1 recognition and not NAD binding, or whether this region only exists for structural integrity and plays no catalytic or target recognition role at all, cannot be answered without a structure of the Csy complex bound to an AcrIF11 homolog.

Remarkably, an Alphafold3 prediction of the Csy complex and AcrIF11_Pae2_ together (Figure 6A) places D146 and E147, the catalytic residues predicted to stabilize the transition state of NAD, in close proximity to N250 of Csy1, the experimentally determined^10^ target residue (Figure 6B). Furthermore, the Alphafold3 prediction suggests that the helical region of AcrIF11_Pae2_ is important for target recognition: residues K58 and K60 of Csy3, residues that have been experimentally confirmed to be important for AcrIF11_Pae1_ activity^10^, are in close proximity to negatively charged residues in the alpha helical region of AcrIF11_Pae2_, revealing a possible charged interaction interface. Likewise, Csy3 residues that were previously determined to have a weaker effect on AcrIF11_Pae1_ activity upon mutation, K66 and S70, are placed further away from AcrIF11_Pae2_ residues in the Alphafold3 prediction (Figure 6C). Because the helical region is poorly conserved across AcrIF11 homologs, we hypothesize that this region confers specificity of an AcrIF11 homolog to a particular target Csy ortholog, or more broadly facilitates these homologs’ ability to distinguish between different Csy orthologs. The structural basis for the high specificity of AcrIF11_Pae2_ that we observed in our lysate blots remains to be explored.

While the enzymatic mechanism of AcrIF11 appears highly potent and specific, enzymatic Acr proteins remain scarce in the literature. This likely represents a discovery challenge that may be solved in the future by integrating structure prediction for *in silico* screening^32^ of common enzymatic folds. It is additionally possible that enzymatic Acrs are rarer for one of two potential reasons: first, the evolution of Acr activity from an enzyme capable of a specific chemistry is likely to happen less often than Acr activity arising from any given non-enzyme. Second, there could be a mechanistic drawback to covalent modifications, as the modification could be undone by host enzymes. The reversal of ADP-ribosylation is an emerging theme in toxin/antitoxin systems important for bacterial and phage ecology^33^. While we could not identify an endogenous bacterial macrodomain that reverses the modification of Csy1, in this study we have established it is at least possible to remove ADP-ribose from Csy1 using human MacroD2. Furthermore, it would make sense for bacteria to have evolved a way to undo enzymatic modifications from phages, especially if the modifications are on a critical residue, such as N250 of Csy1, which functions in PAM recognition. In this context, an enzymatic Acr’s effectiveness would be limited by its high specificity. Stable stoichiometric Acrs, on the other hand, would not be so easily counteracted by the host because they have a larger binding interface with the CRISPR-Cas complex^34^. We envision host-encoded proteins that undo Acr-induced PTMs or PTMs from other phage modification enzymes (e.g. the ART Mod from phage T4)^35^ as a new area to explore in host-phage biology.

## Methods

### AcrIF11_Pae2_ protein purification

#### pET28b-AcrIF11_Pae2_ vector

AcrIF11_Pae2_ coding sequence MGSSHHHHHHSSGLVPRGSHMASMTGGQQMGRMEIFHTSPVEITTINTQGRFGEFLCFAADEYVMTAGDHVTYRIKVDESDIIMAGSIFYHERAADLSGLVERVMQLTGCDEDTAEELISQRIDVFNLDDIDASDAAELSWEIQAITAKAAKTLGFRGVSMQDEQGTCYMIDMLGHDAELVRVK*

AcrIF11_Pae2_ sequence after Thrombin cleavage GSHMASMTGGQQMGRMEIFHTSPVEITTINTQGRFGEFLCFAADEYVMTAGDHVTYRIKVDESDIIMAGSIFYHERAADLSGLVERVMQLTGCDEDTAEELISQRIDVFNLDDIDASDAAELSWEIQAITAKAAKTLGFRGVSMQDEQGTCYMIDMLGHDAELVRVK*

#### 6x His-tag

Thrombin cleavage site

AcrIF11_Pae2_ accession: WP_033936089.1

#### Transformation protocol

BL21(DE3) E. coli were transformed as follows. Frozen stocks were thawed on ice. Upon thawing, 100 ng of the relevant plasmid was added. After a 30 min incubation on ice, cells were heat-shocked for 30 sec at 42 °C and allowed to recover on ice for 2 min. Following, 350 μL of Super Optimal broth with Catabolite repression (S.O.C.) was immediately added and cells were recovered at 37 °C shaking for 1 hour before plating 50 µL on LB agar plates containing appropriate antibiotics. Plates were incubated overnight at 37 °C.

#### M9 Minimal Media Expression for labeled protein expression

Starter cultures of 20 mL of Miller’s LB Broth supplemented with 50 µg/mL kanamycin were inoculated with a single colony of BL21(DE3) E. coli cells harboring pET28b-AcrIF11_Pae2_ and grown overnight for 16 h at 37 °C with shaking at 220 rpm until the culture was saturated. 8 mL of the starter culture was harvested by centrifugation at 4000 x g at 4 °C for 15 minutes. The supernatant was discarded and the pellet was gently resuspended in 5 mL 1X M9 salts. The 5 mL inoculum was then transferred into 1L of M9 minimal media containing appropriate labeling components (see M9 minimal media assembly) and 50 µg/mL kanamycin. Cells were then grown at 37 °C with shaking at 220 rpm to an OD of 1.0 at which point the culture was induced with 1mL of 1 mM IPTG and grown for 16 hours at 16 °C. The expression culture was harvested by centrifugation at 5000 x g at 4 °C for 20 minutes. Pellets were either stored at −80 °C or immediately used for purification.

#### Protein purification

The resulting cell pellet was suspended in 20 mL of Buffer A (30 mM Imidazole, 250 mM NaCl, 20 mM HEPES pH 7.0, 5% Glycerol, and 0.5 mM TCEP) containing a tablet of cOmplete, mini EDTA-free ULTRA protease inhibitor cocktail. The cell suspension was disrupted by sonication on ice (1 second pulse at 50% duty cycle followed by 1 second pause for a total time of 5 min). The cell lysate was cleared by centrifugation at 30,000xg at 4 °C for 30 minutes. Nickel affinity purification was conducted with a Cytiva 5 mL HisTrap HP column. The column was first equilibrated with Buffer A over 5 column volumes (CV), then the lysed sample was applied to the column at a flow rate of 3 mL/min. After protein was bound to the column, a wash was performed with 95% Buffer A and 5% Buffer B (500 mM Imidazole, 250 mM NaCl, 20 mM HEPES pH 7.0, 5% Glycerol, and 0.5 mM TCEP) over 10 CV. The protein was eluted over a linear gradient from 95% Buffer A and 5% Buffer B to 5% Buffer A and 95% Buffer B over 5 CV and fractionated. Fractions containing protein were concentrated using a Amicon^®^ Ultra Centrifugal Filter 10 kDa MWCO to 10 mL. 10 units/mg thrombin was added to the protein, and the mixture was then dialyzed into Buffer C (100 mM NaCl, 20 mM HEPES pH 7.0, and 0.5 mM TCEP) at 4 °C overnight. The following day, a reverse nickel affinity purification was run using the same method as nickel affinity purification, but with replacing Buffer A with Buffer C. Fractions were collected as the sample was applied to the column to capture cleaved AcrIF11_Pae2_ protein. AcrIF11_Pae2_ protein fractions were concentrated to 2 mL and immediately applied to a HiLoad^®^ 16/600 Superdex^®^ 75 pg SEC column using Buffer C over 1.3 CV and fractionated. Fractions containing the protein were concentrated to 3 mL and then desalted using a Bio-Gel p-6DG gel Desalting column in accordance with manufacturer instructions. The purified H3 protein was quantified using absorbance at 280 nm and concentrated to a final protein concentration of 1 mM for NMR analysis.

#### M9 minimal media assembly

M9 media was prepared accordingly for either ^15^N-labeled AcrIF11_Pae2_ protein expression or ^15^N-/^13^C-labeled H3 protein expression (Supplementary Table 3 and 4). First, 10x M9 salts, ammonium sulfate, and MilliQ H2O were autoclaved together. The remaining components were sterile filtered individually into the autoclaved solution to create 1 L of M9 media for labeled protein expression.

### AcrIF11_Pae2_ NMR structure determination

The final protein concentration for the [U^13^C, U^15^N]-labeled protein for the data collection for the structure determination^36^ was 1mM AcrIF11_Pae2_ in 50 mM KPi pH 7.4 with 5% (v/v) D2O. NMR spectra were all measured at 298.0 K^37^. Two dimensional (2D) 1H,15N-HSQC (pulse program: fhsqcf3gpph), 2D multiplicity-edited CT 1H,13C-HSQC (pulse program: hsqcctetgpsp), 36ms 3D HCcH-TOCSY (pulse program hcchdigp3d) and 120ms 3D simultaneous 13C-/15N-NOESY-HSQC (pulse program: noesyhsqcgpsismsp3d) spectra were measured using a Bruker Avance NEO 800 MHz spectrometer with a 5mm TCI H&F-C/N-D-05 Z-gradient CP2.1 CryoProbe. 3D CACBcoNH (pulse program: cbcaconhgp3d), 3D HNCACB (pulse program: hncacbgp3d), 3D HcccoNH (pulse program: hccconhgp3d2), 3D hCccoNH (pulse program: hccconhgp3d3) experiments were collected on a Bruker Avance NEO 600 MHz spectrometer with an 5mm TCI H&F-C/N-D-05 Z-gradient CP2.1 CryoProbe. Spectra were processed in TopSpin version 4.0.6 and referenced indirectly to an external DSS standard^38^.

Resonances were assigned using the program CCPN Analysis version 2.4.2^39^. Backbone resonances were assigned using the 2D 15N-HSQC, 3D CACBcoNH and 3D HNCACB spectra. Sidechain resonances were assigned using the 2D constant-time 13C-HSQC and 3D HCcH-TOCSY spectra. Distance restraints were generated using CCPN Analysis. Dihedral restraints were generated using the program DANGLE^40^. The programs ARIA version 2.3.2^41^ and CNS version 1.2.1^42^ were used to calculate the NMR structures. To solve the structures, 9 iterations of simulated annealing were performed using CNS. For the first 8 rounds of simulated annealing, the n_structures parameter was set to 50 and the n_best_structures parameter was set to 15. For the 8th round, the n_structures parameter was set to 200 and the n_best_structures parameter was set to 25. Finally, a refinement in water was performed on the lowest energy structures from the 8th iteration. Otherwise, the default values were used for the remaining ARIA parameters. Initial structure calculations were conducted without hydrogen bond restraints. Hydrogen bond donors were then identified, and the corresponding hydrogen bond restraints were included in later calculations. Hydrogen bond restraints included in the structure calculations were based on measurements of amide chemical exchange with solvent detected by 2D ^15^N-CLEANEX-HSQC experiments^43^. Structures were validated using the Protein Structure Validation Software (PSVS) suite 1.5. The chemical shifts, restraints, and structural coordinates have been deposited with the BMRB (31035) and PDB (8DWQ).

### AcrIF11 phylogenetic analysis

To find AcrIF11 homologs, AcrIF11_Pae1_ (WP_038819808.1) was PSI-BLAST was used with the nr database for 3 iterations. Hits with greater than 70% coverage and expected value less than 0.0005 were used for generating the PSSM in each iteration, and the maximum hitlist size was set to 500. Sequences were aligned via MAFFT^44^, and the multiple sequence alignment was trimmed manually to prevent large gaps in the alignment. Sequences larger than 200 residues were also discarded to prevent large gaps in the alignment. Iterative rounds of trimming and alignment were performed to refine the alignment until gaps were smaller than 25 residues. The phylogenetic tree was created from the final multiple sequence alignment using FastTree^45^, and visualization was done using iTOL^46^.

For determining the distribution of Type I-F Acrs, the Type I-F Acr sequences were first acquired from the Acr database^47^. The sequences were blasted against the clustered_nr database using BLASTp, with an expectation value of 0.0005 and a maximum hitlist size of 5000. Unique genera were manually counted from the hitlists.

### Alphafold structure prediction

The hits used to make the phylogeny in “AcrIF11 phylogenetic analysis” were clustered using MMseqs2^48^, using minimum sequence identity of 95%, cluster mode 2, and coverage mode 1. Sequences were also trimmed for length and to prevent large gaps in alignment. Then the representative sequences of each cluster were used for structure prediction. Sequences were run through the SBGrid installation of Alphafold2, with a maximum template date set to 2020-09-20. The highest confidence results (structures named ranked_0.pdb) were used for sequence conservation and RMSD analysis via MatchMaker in ChimeraX^49^.

The Alphafold3 web server (https://alphafoldserver.com/) was used to predict AcrIF11 with the Csy complex. Sequences used for Alphafold3 prediction were from Guo et. al^50^ PDB 6B45: AcrIF11_Pae2_: MEIFHTSPVEITTINTQGRFGEFLCFAADEYVMTAGDHVTYRIKVDESDIIMAGSIFYHE RAADLSGLVERVMQLTGCDEDTAEELISQRIDVFNLDDIDASDAAELSWEIQAITAKAAK TLGFRGVSMQDEQGTCYMIDMLGHDAELVRVK

Csy1: MTSPLPTPTWQELRQFIESFIQERLQGKLDKLQPDEDDKRQTLLATHRREAWLADAARRV GQLQLVTHTLKPIHPDARGSNLHSLPQAPGQPGLAGSHELGDRLVSDVVGNAAALDVFKF LSLQYQGKNLLNWLTEDSAEALQALSDNAEQAREWRQAFIGITTVKGAPASHSLAKQLYF PLPGSGYHLLAPLFPTSLVHHVHALLREARFGDAAKAAREARSRQESWPHGFSEYPNLAI QKFGGTKPQNISQLNNERRGENWLLPSLPPNWQRQNVNAPMRHSSVFEHDFGRTPEVSRL TRTLQRFLAKTVHNNLAIRQRRAQLVAQICDEALQYAARLRELEPGWSATPGCQLHDAEQ LWLDPLRAQTDETFLQRRLRGDWPAEVGNRFANWLNRAVSSDSQILGSPEAAQWSQELSK ELTMFKEILEDERD

Csy2: MSVTDPEALLLLPRLSIQNANAISSPLTWGFPSPGAFTGFVHALQRRVGISLDIELDGVG IVCHRFEAQISQPAGKRTKVFNLTRNPLNRDGSTAAIVEEGRAHLEVSLLLGVHGDGLDD HPAQEIARQVQEQAGAMRLAGGSILPWCNERFPAPNAELLMLGGSDEQRRKNQRRLTRRL LPGFALVSREALLQQHLETLRTTLPEATTLDALLDLCRINFEPPATSSEEEASPPDAAWQ VRDKPGWLVPIPAGYNALSPLYLPGEVRNARDRETPLRFVENLFGLGEWLSPHRVAALSD LLWYHHAEPDKGLYRWSTPRFVEHAIA

Csy3: MSKPILSTASVLAFERKLDPSDALMSAGAWAQRDASQEWPAVTVREKSVRGTISNRLKTK DRDPAKLDASIQSPNLQTVDVANLPSDADTLKVRFTLRVLGGAGTPSACNDAAYRDKLLQ TVATYVNDQGFAELARRYAHNLANARFLWRNRVGAEAVEVRINHIRQGEVARAWRFDALA IGLRDFKADAELDALAELIASGLSGSGHVLLEVVAFARIGDGQEVFPSQELILDKGDKKG QKSKTLYSVRDAAAIHSQKIGNALRTIDTWYPDEDGLGPIAVEPYGSVTSQGKAYRQPKQ KLDFYTLLDNWVLRDEAPAVEQQHYVIANLIRGGVFGEAEEK

Csy4: MDHYLDIRLRPDPEFPPAQLMSVLFGKLHQALVAQGGDRIGVSFPDLDESRSRLGERLRI HASADDLRALLARPWLEGLRDHLQFGEPAVVPHPTPYRQVSRVQAKSNPERLRRRLMRRH DLSEEEARKRIPDTVARALDLPFVTLRSQSTGQHFRLFIRHGPLQVTAEEGGFTCYGLSK GGFVPWF

crRNA: CUAAGAAAUUCACGGCGGGCUUGAUGUCCGCGUCUACCUGGUUCACUGCCGUGUAGGC AG

### Determining the presence of AcrIF11 on mobile genetic elements

The first mode of analysis consisted of blasting AcrIF11 homologs from the phylogenetic analysis above (only the representative sequences of each cluster) against a homemade database of plasmid and phage proteins. Plasmid proteins were downloaded from the NCBI RefSeq Plasmid database (https://ftp.ncbi.nlm.nih.gov/genomes/refseq/plasmid/), and phage proteins were downloaded from the NCBI Virus web database (Find Data > Search by virus > Bacteriophages). The BLAST database combining the plasmid and phage dataset was created using makeblastdb in the BLAST command line application. AcrIF11 homologs were blasted against this database using BLASTp with an expectation value of 0.0005. Results were marked as “phage” if the species indicated in the hitlist (or species associated with the query accession) was a phage, and “plasmid” if the species indicated in the hitlist was a bacterium. Hits were only considered valid if the query and target length matched exactly, and if the sequence identity was 100% with zero gaps.

The second mode of analysis was using geNomad^16^. For every AcrIF11 homolog, the associated genome accessions were identified using eLink from NCBI eUtilities, or manually in cases where eLink failed. geNomad was run on each genome, with score calibration enabled and post-classification filtering set to conservative. The AcrIF11 homologs associated with each genome were then blasted against said genome to identify its genomic location. The genomic location of the AcrIF11 homolog was then compared to the prophage location identified by geNomad, or to the plasmid genes location list identified by geNomad, to ensure that each homolog fell within the bounds of the mobile genetic element. In the case of plasmid hits, which were identified by coding region accession numbers instead of translated protein accession numbers, the hits were also confirmed to be the homolog of interest via NCBI Identical Protein Groups. Results appearing as “prophage + plasmid” either had both hits in geNomad, or a hit in geNomad (prophage) and a hit in the homemade database (plasmid).

### CRISPRi pyocyanin assay

This assay was performed as previously described^24^. Briefly, a plasmid encoding a Type I-F crRNA targeting the *phzM* (pyocyanin synthesis gene) promoter was used to transform the desired lysogen. An empty vector was also transformed into the lysogen as a control. Lysogens were grown as overnight cultures with gentamicin for plasmid maintenance and 0.1% arabinose for induction of crRNA expression. Pyocyanin was extracted with an equal volume of chloroform, mixed with a half volume of 0.2M HCl, and quantified by measuring absorbance at 520 nm. For the plot in Fig 2C, pyocyanin levels were normalized to the empty vector control.

### Molecular cloning & PA14 conjugation

Genes of interest were synthesized or PCRed from template DNA, and then inserted into pHERD20T or pHERD30T backbone using the NEB-recommended HiFi protocol. pHERD vectors were digested with NcoI-HF and Sbf-HF (for 20T) or NcoI-HF and HindIII-HF (for 30T) for the HiFi reaction. XL1-B cells were transformed with the HiFi reactions, and transformants were verified via Sanger sequencing. Miniprepped plasmids were verified again via whole-plasmid sequencing from Primordium.

To introduce plasmids into PA14, *E. coli* SM10 cells were used as donors. Plasmids were electroporated into SM10 cells and plated onto LB + antibiotic plates (carbenicillin for 20T, gentamicin for 30T) after recovering in LB at 37°C for 2-3 hours. SM10 transformed colonies were cross-streaked with PA14 for conjugation. After incubating conjugation plates overnight at 37°C, PA14 colonies that received the plasmid were selected via VBMM + antibiotic plates. Selected colonies were verified via PCR and Sanger sequencing of PCR products.

### Liquid phage infections

PA14 strains were grown in LB or LB + the appropriate antibiotic overnight at 37°C and diluted 1:100 in the same LB formulation. 10-fold serial dilutions of the desired phage were made using SM buffer. In each well of a 96-well plate, 10 µL of each phage dilution was added to 140 µL of 1:100 diluted overnight culture, and grown at 37°C on a plate reader that monitored OD600.

### Plaque assays

PA14 strains were grown in LB or LB + the appropriate antibiotic overnight at 37°C. 150 µL of the overnight culture was mixed with 3 mL of LB top agar supplemented with 10 mM Mg (and arabinose at 0.1% when working with a strain containing a pHERD vector). The top agar + overnight culture mixture was plated onto LB + 10 mM Mg + appropriate antibiotic plates. 10-fold serial dilutions of the desired phage were made using SM buffer, and 2.5 µL of each dilution was spotted onto the plate. Plates were incubated at 30°C overnight.

### Full plate phage infections

PA14 strains were grown in LB or LB + the appropriate antibiotic overnight at 37°C. 150 µL of overnight culture was added to 10 µL of phage, and mixed via shaking at room temperature for 15 minutes. This mixture was then added to 3 mL of LB top agar supplemented with 10 mM Mg, and spread out over an LB + 10 mM Mg + appropriate antibiotic plate. Plates were incubated at 30°C overnight. Plaques were manually counted. Phage titers were calculated via full plate infections with a PA14 Δcr1Δcr2 lawn.

### Phage competition

PA14 WT, containing either Csy1-4 on pHERD30T or empty vector pHERD30T, was grown in LB + 10 mM Mg + Gent50 overnight at 37°C and diluted 1:100 in LB + 10 mM Mg + Gent50 + 0.1% arabinose. DMS3mF11_Pae1_vir and DMS3mF1vir were diluted to the appropriate concentration in SM buffer. In each well of a 96-well plate, 140 µL of the 1:100 diluted overnight culture was added, and either 10 µL of one phage was added for single-phage growths, or 5 µL of each phage for mixed phage growths. Plates were grown at 37°C on a plate reader that monitored OD600. After the growth period was completed, phages were extracted from plate reader cultures using 6-8% chloroform. These extracted phages were used in full plate infections with PA14 Δcr1Δcr2, with each MOI or MOI combination indicated in Fig 3E on a separate plate. 16 plaques were picked for each MOI or MOI combination, and subject to PCR using primers surrounding DMS3 gene 30 (where the appropriate *acr* gene was inserted). Based on the size of the PCR product viewed on an agarose gel, the identity of the Acr was determined (AcrIF1 is approximately 300 bp, while AcrIF11_Pae1_ is approximately 500 bp).

### Lysogen construction

Plaque assays with spot titrations of the desired phage were done following the protocol outlined above. Clearings from plaque assays were streaked out onto LB + 10 mM Mg plates and incubated overnight at 37°C. The resulting colonies were grown up as overnight cultures in LB + 10 mM Mg, and lysogeny was verified via two methods: plating the putative lysogen culture and spotting with the same phage to check for superinfection exclusion, and chloroform extraction of the putative lysogen culture followed by spotting the extract onto a PA14 lawn to check for the presence of virions.

### Lysogen growth experiments

Lysogens were streaked out onto LB + 10 mM Mg plates from glycerol stocks. LB + 10 mM Mg cultures were inoculated with a lysogen colony and grown overnight at 37°C. The overnight culture was passaged 1:100 into LB+Mg three times, with the OD600 of each passage monitored via plate reader for 20 hours. Three replicates were performed, with each replicate consisting of three passages.

### Western blot of lysates

Three replicates of the AcrIF11_Pae2_ mutant blots were done. For each replicate, the desired plasmid was conjugated into PA14 sfCherry2-Csy1 following the conjugation protocol outlined above. Conjugated colonies were verified via PCR of the multiple cloning site and Sanger sequencing, then grown overnight in LB + Carb250. Overnight cultures were diluted 1:100 in fresh LB + Carb250 + 0.1% arabinose, and grown for 13 hours at 37°C. After 13 hours, cultures were pelleted, washed, resuspended in lysis buffer, incubated in lysis buffer for 15 minutes, and then lysed via sonication with the Bioruptor Pico. The wash buffer used was 50 mM Tris pH 7.4, 150 mM NaCl, 1 mM EDTA, 1 mM MgCl_2_. The lysis buffer contained the same formulation as the wash buffer, but with the addition of 1 mg/mL lysozyme, 1 protease inhibitor tablet, 0.5 mM TCEP, and 125 U/mL Pierce Universal Nuclease. After lysis, lysates were clarified with a 20 minute spin in a benchtop centrifuge at 4°C and approximately 21,000 x g.Clarified lysates were then flash frozen in liquid nitrogen for future blotting. For lysogen lysate blots, three replicates were also grown and lysed in the same manner as above, but grown with LB + 10 mM Mg instead of LB + Carb250 + 0.1% arabinose.

Total protein concentration in lysate was quantified via Bradford assay. Lysates were prepared for SDS-PAGE by resuspending in SDS loading buffer and heating at 95°C for 5 minutes. Gel samples were loaded onto an SDS-PAGE gel, with loading volumes adjusted to normalize total protein concentration across all wells. Depending on the particular concentrations from lysis in each replicate, 15-30 µg/well was loaded. Gels were transferred onto a nitrocellulose membrane using the BioRad TurboBlot, and normalization was verified via Ponceau staining. After washing off Ponceau stain, blots were blocked for 1 hour at room temperature in 5% casein, and then incubated in 1:1000 of primary antibody (ADPr: Cell Signaling D9P7Z, Cherry: Cell Signaling E5D8F) overnight at 4°C. The next day, the blots were washed in TBS-T for 10 minutes per round, for 3 rounds. Blots were blocked in 1:10,000 HRP secondary antibody (Cell Signaling 7074) at room temperature for 1 hour, then washed in TBS-T for 10 minutes per round, for 3 rounds. BioRad Clarity Max ECL substrate was added to the blots in accordance with the manufacturer protocol, and blots were imaged using the BioRad Chemidoc. Blots for ADPr and Cherry were run in parallel with the same lysates and same loading volumes. During initial trial runs of the ADPr blots, we noticed significant differences in signal detection between ADPr antibodies from different manufacturers, as well as sample preparation and storage conditions. This is in line with previous observations of ADPr reagent variability^51^, and for consistency we decided to use only the Cell Signaling primary mentioned above.

### Western blot quantification

Image Lab 6.1 from Bio-Rad was used for quantification of band intensity in Fig 5B. In the Analysis Toolbox, lanes and the appropriate bands were selected using the tools in the Lanes and Bands menu. Background subtraction and accurate band selection were manually verified using the Lane Profile tool. Adjusted volume of each band was normalized to the adjusted volume of the +AcrIF11_Pae2_, -macrodomain band (leftmost lane of Fig 5B).

### hMacroD2 macrodomain protein purification

#### hMacroD2 sequence

MHHHHHHSSGVDLGTENLYFQSYPSNKKKKVWREEKERLLKMTLEERRKEYLRDYIPLNSILSWKEEMKGKGQNDEENTQETSQVKKSLTEKVSLYRGDITLLEVDAIVNAANASLLGGGGVDGCIHRAAGPCLLAECRNLNGCDTGHAKITCGYDLPAKYVIHTVGPIARGHINGSHKEDLANCYKSSLKLVKENNIRSVAFPCISTGIYGFPNEPAAVIALNTIKEWLAKNHHEVDRIIFCVFLEVDFKIYKKKMNEFFSVDDNNEEEEDVEMKEDSDENGPEEKQSVEEMEEQSQDADGVNTVTVPGPASEEAVEDCKDEDFAKDENITKGGEVTDHSVRDQDHPDGQENDSTKNEIKIETESQSSYMETEELSSNQEDAVIVEQPEVIPLTEDQEEKEGEKAPGEDTPRMPGKSEGSSDLENTPGPDAGAQDEAKEQRNGTKGLNDIFEAQKIEWHE*

#### 6xHIS / AVI / TEV

hMacroD2 was expressed and purified as described previously for SARS-CoV-2 Mac1^52^.

### BFV macrodomain protein purification

#### BFV sequence

MSYYHHHHHHLESTSLYKKAGFLEVLFQGPEVNSFSGYLKLAPAYRVKRGDISNAPEDAVVNAANQQGVKGAGVCGAIYRKWPDAFGDVATPTGTAVSKSVQDKLVIHAVGPNFSKCSEEEGDRDLASAYRAAAEIVMDKKITTVAVPLLSTGIYAGGKNRVEQSLNHLFTAFDNTDADVTIYCMDKTWEKKIKEAIDHRT

#### Cloning and expression

The macrodomain of BFV was synthesized and cloned into the pET28 vector, incorporating an N-terminal 6xHis tag and a long linker before the macrodomain, resulting in the expression construct 6xHis-LINKER-Macrodomain. The plasmid was then transformed into LOBSTR-BL21(DE3) *E. coli* for protein expression. Following a standard expression protocol, cells were grown in LB medium at 37 °C until the OD at 600 nm reached 0.8. Protein expression was then induced by the addition of 500 µM IPTG, and the cells were allowed to grow overnight at 18 °C. After incubation, the cells were harvested by centrifugation. The cell pellets were recovered and stored at −80 °C until purification.

#### Protein purification

The cell pellet was suspended in lysis buffer (50 mM Tris, pH 7.0, 250 mM NaCl, 10 mM imidazole, 5% glycerol, and 2 mM β-mercaptoethanol) containing one tablet of Complete, Mini EDTA-free ULTRA protease inhibitor cocktail and DNase I (10 µg/mL). The cell suspension was disrupted by passaging three times through a chilled Emulsiflex at 15,000 psi. The cell lysate was clarified by centrifugation at 30,000 × g for 30 minutes at 4 °C.

Nickel affinity purification was conducted using a gravity flow column packed with 5 mL of resin. The column was first equilibrated with water and then with lysis buffer. Following equilibration, the lysate was added to the resin and rocked at 4 °C for 30 minutes. After the protein bound to the column, the column was washed with wash buffer (50 mM Tris, pH 7.0, 250 mM NaCl, 10 mM imidazole, 5% glycerol, and 2 mM β-mercaptoethanol) over 2 × 10 column volumes (CV). The protein was eluted with elution buffer (50 mM Tris, pH 7.0, 250 mM NaCl, 500 mM imidazole, 5% glycerol, and 2 mM β-mercaptoethanol) in 6 × 5 mL fractions (30 mL total).

Fractions containing the protein were pooled and dialyzed overnight at 4 °C to remove imidazole using a dialysis buffer (50 mM Tris, 150 mM NaCl, 1 mM DTT, 5% glycerol). The protein was concentrated to 1 mL and immediately applied to a Superdex® 75 10/300 SEC column using size exclusion buffer (50 mM Tris, pH 7.0, 150 mM NaCl, 5% glycerol). Fractions containing the protein were concentrated to 0.7 mg/mL, flash-frozen, and stored at −80 °C until required for assays.

### EEEV macrodomain protein purification

#### EEEV sequence

MGHHHHHHHHHHENLYFQSGAPAYRVVRGDITKSNDEVIVNAANNKGQPGGGVCGALYRKWPGAFDKQPVATGKAHLVKHSPNVIHAVGPNFSRLSENEGDQKLSEVYMDIARIINNERFTKVSIPLLSTGIYAGGKDRVMQSLNHLFTAMDTTDADITIYCLDKQWESRIKEAI

#### Cloning and expression

The macrodomain of EEEV was synthesized and cloned into the pET28 vector, incorporating an N-terminal 6X-HIS tag, TEV protease site and linker before macrodomain, generating the expression as 6XHis-TEV-SG-Macrodomain. The plasmid was then transformed into LOBSTR-BL21(DE3) E. coli for protein expression. Following a standard expression protocol, cells were grown in LB media at 37°C until OD 600 nm reached 0.8, then protein expression was induced by the addition of 500 μM IPTG to the media, and cells were allowed to grow overnight at 18°C, after which they were harvested by centrifugation. Cell pellets were recovered and stored at −80°C until purification.

#### Protein purification

Purifications were carried out under standard NTA purification conditions. Briefly, cells pellets were resuspended in lysis buffer, 50 mM Tris pH 8, 500 mM NaCl, 10mM imidazole, 5% glycerol, 2 mM β-mercaptoethanol. Cells were then lysed by passing through chilled Emulsiflex at ∼10,000psi for three cycles. The lysate was clarified by centrifugation at 35,000 g for 1 hour. The clarified lysate was applied to 5 ml of pre-equilibrated Ni-NTA resin and incubated for 1 hour. The resin was then washed with 5 times 10 CV of 50 mM Tris pH 8, 500 mM NaCl, 30mM imidazole, 5% glycerol, 2 mM β-mercaptoethanol. Elution of the protein was done with 5 CV buffer 50 mM Tris pH 8, 500 mM NaCl, 500 mM imidazole, 5% glycerol, 2mM BME. The eluted protein was dialyzed in 50LJmM Tris, pH 8.0, 150LJmM NaCl, 5% glycerol, 1LJmM DTT. The His-tag was cleaved with purified His-TEV protease (1:20 mass ratio with EEEV eluted protein). His-TEV was further removed by reverse NTA purification. Further EEEV macrodomain was purified using SEC S75 10/300 column in 20LJmM Tris, pH 7, 150LJmM NaCl and fractions with the purest protein were mixed, concentrated, flash frozen and stored at −80°C until required for assays.

### Macrodomain reactions

AcrIF11_Pae2_ WT was conjugated into PA14 sfCherry2-Csy1, grown, and lysed following the same protocol as outlined above in “Western blot of lysates”. Before loading onto the SDS-PAGE gel, purified macrodomain was added and incubated with lysates for 1 hour at room temperature. SDS loading buffer was added to the reactions, and heated for 5 minutes at 95°C before running the SDS-PAGE gel. The blotting procedure was the same as outlined above, but after imaging with the ADPr primary antibody, the blot was stripped using Restore Plus Stripping Buffer for 10 minutes at 37°C, blocked using the same protocol as above, and incubated overnight in Cherry primary antibody solution. The next day, the same protocol as above was done for washing, secondary incubation, and imaging.

### Phage genetic editing

*acrIF1* naturally overlaps with a downstream gene *aca1*, a transcriptional regulator of Acr expression. The native architecture of this operon was maintained when introducing *acrIF1* into DMS3m/DMS3mvir phages. *acrIF11_Pae1_*does not co-occur with *aca1* and thus was engineered in with *aca1* downstream, but no overlap. These genes were engineered into DMS3m/DMS3mvir phages as previously described.^19^ Briefly, each gene was introduced into the pHERD plasmid with homology regions flanking DMS3 gene 30. Cells containing the plasmid were infected with DMS3m/DMS3mvir, and phages isolated from this infection were then plated onto PA14, which contains a Type I-F CRISPR system to select for recombinant DMS3m/DMS3mvir. Individual plaques from this selection were purified and the presence of the *acr* was verified via Sanger sequencing.

## Supplementary Information

**Supplementary Table 1.** NMR assignment, structure calculation and validation statistics.

|  |  |
| --- | --- |
| **Degree of assignment<sup>a</sup>** |  |
| Backbone (N and H<sup>N</sup>) (%) | 89.8 |
| Side-chain H (%) | 72.4 |
| Side-chain non-H (%) | 56.9 |
| **Number of restraints<sup>b</sup>** |  |
| NOE restraints |  |
| Intra-residue ($\vert i-j\vert = 0$) | 591 |
| Sequential ($\vert i-j\vert = 1$) | 545 |
| Medium range ($1 < \vert i-j\vert < 5$) | 423 |
| Long range ($\vert i-j\vert \geq 5$) | 537 |
| Ambiguous<sup>a</sup> | 640 |
| Total | 2069 |
| H-bond restraints | 54 |
| Long range ($\vert i - j\vert \geq 5$) | 40 |
| Dihedral angle restraints ($\phi/\psi$) | 150/150 |
| **Restraint statistics<sup>c</sup>** |  |
| r.m.s. of NOE violations (Å) | 0.525 ± 0.498 |
| r.m.s. of dihedral violations (°) | 2.60 ± 2.488 |
| **r.m.s. from idealised covalent geometry<sup>d</sup>** |  |
| Bonds (Å) | 0.0042 ± 0.0002 |
| Angles (°) | 0.63 ± 0.041 |
| Impropers (°) | 2.28 ± 0.24 |
| Structural quality |  |
| Ramachandran statistics<sup>e/f</sup> |  |
| Most favoured regions (%) | 84.4 / 90.2 |
| Allowed regions (%) | 14.9 / 8.5 |
| Generously allowed regions (%) | 0.7 / NA |
| Disallowed regions (%) | 0.0/1.3 |
| Verify3D Z-score<sup>g</sup> | -4.98 |
| Prosa II Z-score<sup>h</sup> | -0.99 |
| Procheck Z-score ($\phi/\psi$)<sup>e</sup> | -1.46 |
| Procheck Z-score (all)<sup>e</sup> | -3.02 |
| MolProbity Z-score<sup>f</sup> | -5.86 |
| No. of close contacts<sup>i</sup> | 11 |
| **Coordinates precision (rmsd)<sup>b</sup>** |  |
| All backbone atoms (Å) | 2.8 / 2.3 |
| All heavy atoms (Å) | 3.3 / 2.8 |

Values reported by: <sup>a</sup>CCPNMR 2.5.2<sup>39</sup>; <sup>b</sup>Protein Structure Validation Software suite 1.5 & <sup>c</sup>PDBStat 5.12<sup>53</sup>; <sup>d</sup>Crystallography and NMR system (CNS) 1.2<sup>54</sup>; <sup>e</sup>Procheck<sup>55</sup> & <sup>f</sup>MolProbity<sup>56</sup>. The structural validation programs used were as follows: <sup>g</sup>Verify3D<sup>57</sup>, <sup>h</sup>Prosa II<sup>58</sup>, <sup>e</sup>Procheck<sup>55</sup>, <sup>f</sup>MolProbity<sup>56</sup> and <sup>i</sup>PDB validation software.

**Supplementary Table 2.**
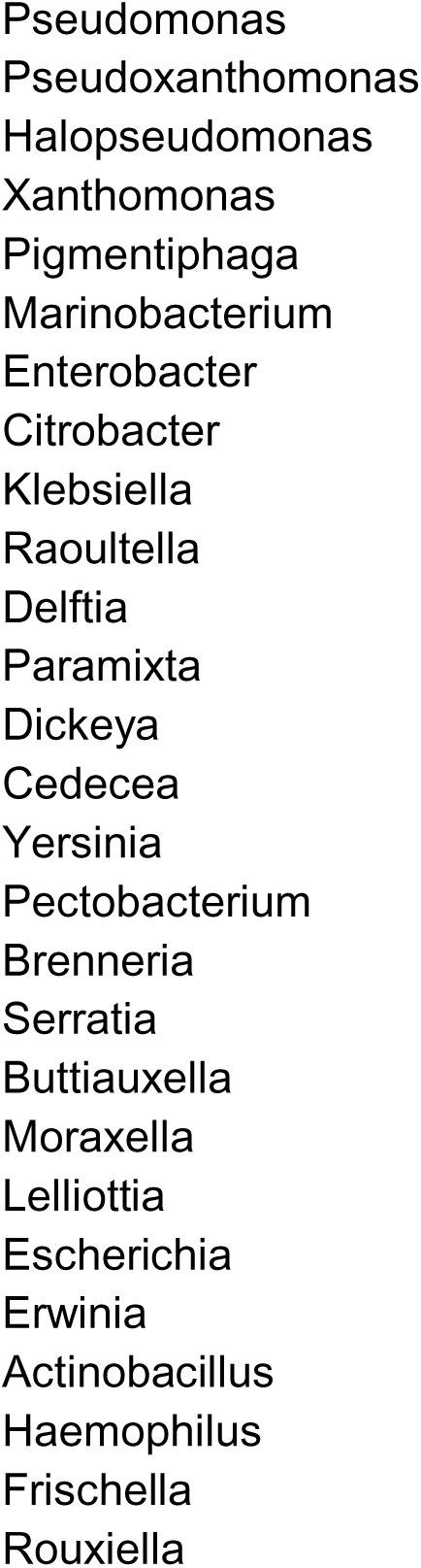
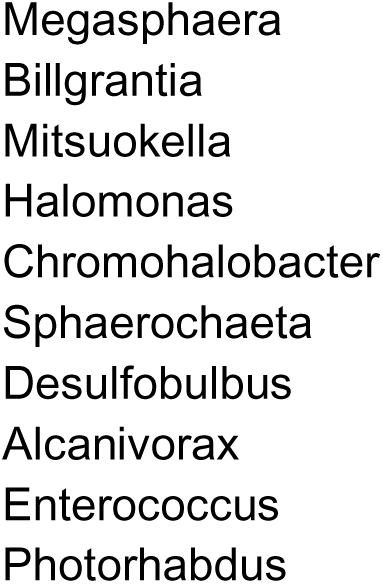
List of bacterial genera containing AcrIF11 homologs.

**Supplementary Table 3.** 1 L M9 Minimal Media Recipe for 15N labeled proteins.

| Component | Amount |
| --- | --- |
| 10x M9 salts | 100 mL |
| 1M Magnesium sulphate | 1 mL |
| 1M Calcium chloride | 100 $\mu\text{L}$ |
| 20% Thiamine | 100 $\mu\text{L}$ |
| 0.003 g/mL Iron II sulphate heptahydrate | 1 mL |
| 10x MEM Vitamin mix | 10 mL |
| 20% Glucose | 40 mL |
| 15N Ammonium sulfate (15N source) | 1 g |
| MilliQ H<sub>2</sub>O | 860 mL |

All solutions are dissolved in MilliQ H<sub>2</sub>O

**Supplementary Table 4.** 1 L M9 Minimal Media Recipe for 15N 13C labeled proteins.

| Component | Amount |
| --- | --- |
| 10x M9 salts | 100 mL |
| 1M Magnesium sulphate | 1 mL |
| 1M Calcium chloride | 100 $\mu\text{L}$ |
| 20% Thiamine | 100 uL |
| Iron II sulphate heptahydrate | 3 mg |
| 10x MEM Vitamin mix | 10 mL |
| 10% 13C Glucose | 40 mL |
| 15N Ammonium sulfate | 1 g |
| MilliQ H2O | 860 mL |

All solutions are dissolved in MilliQ H2O

**Supplementary Figure 1.**
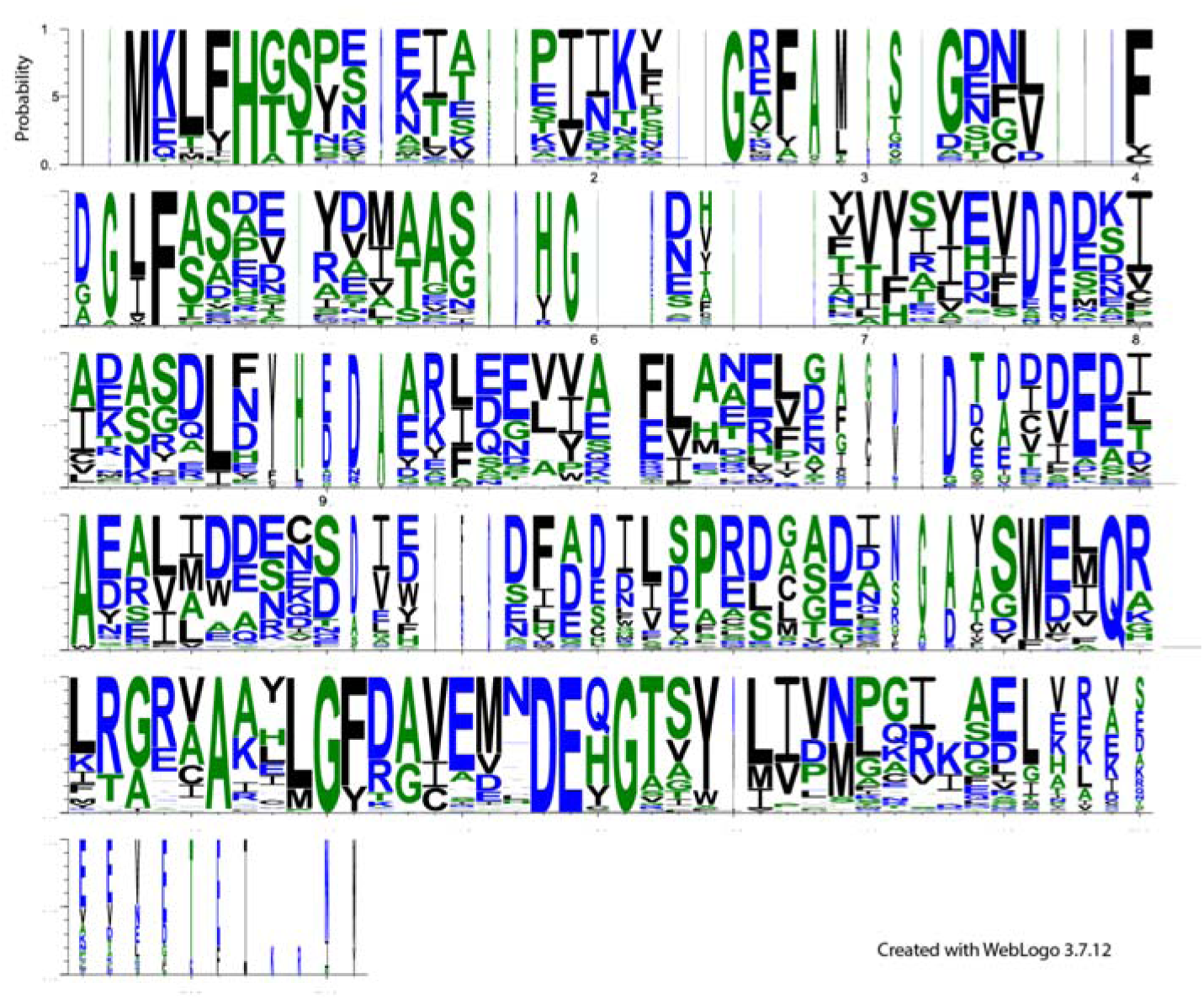
Sequence logo of AcrIF11 homologs.

**Supplementary Figure 2.**
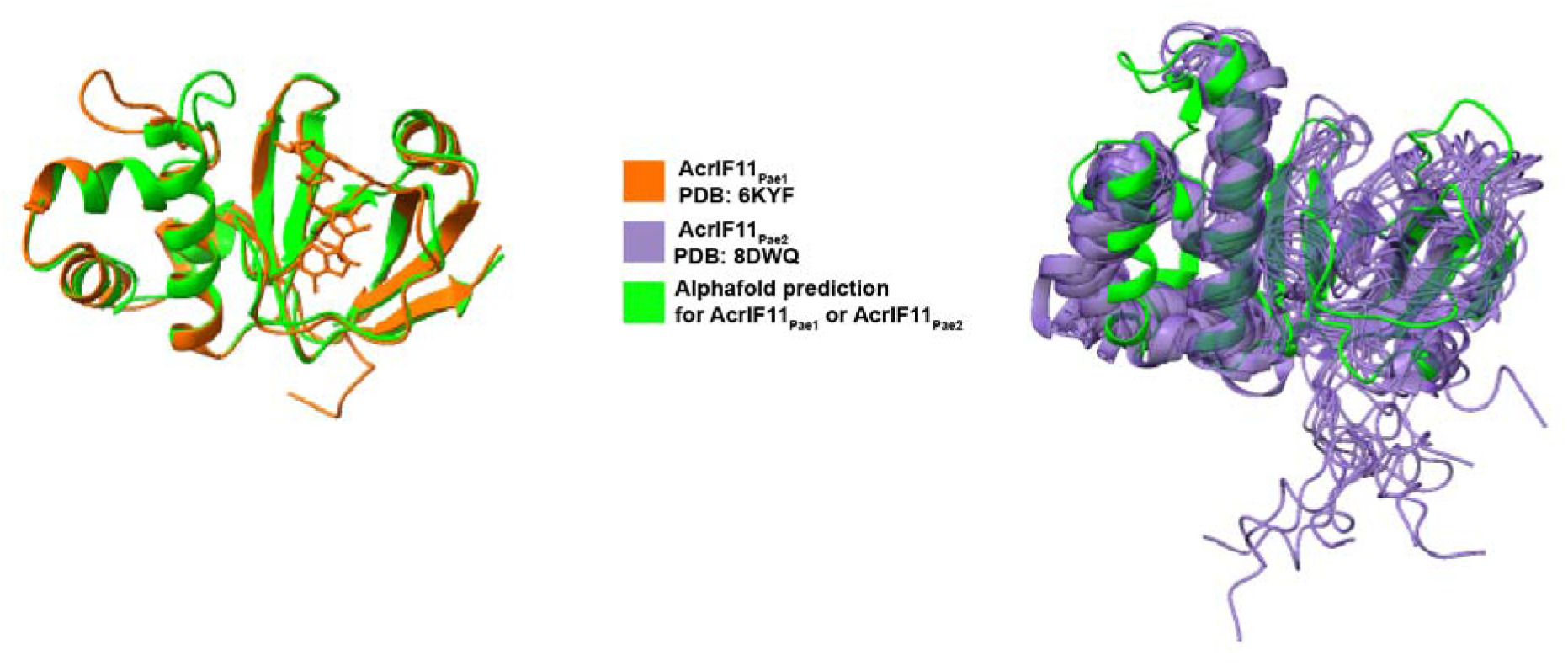
Alignment of experimental and predicted structures of AcrIF11_Pae1_ and AcrIF11_Pae2_

**Supplementary Figure 3.**
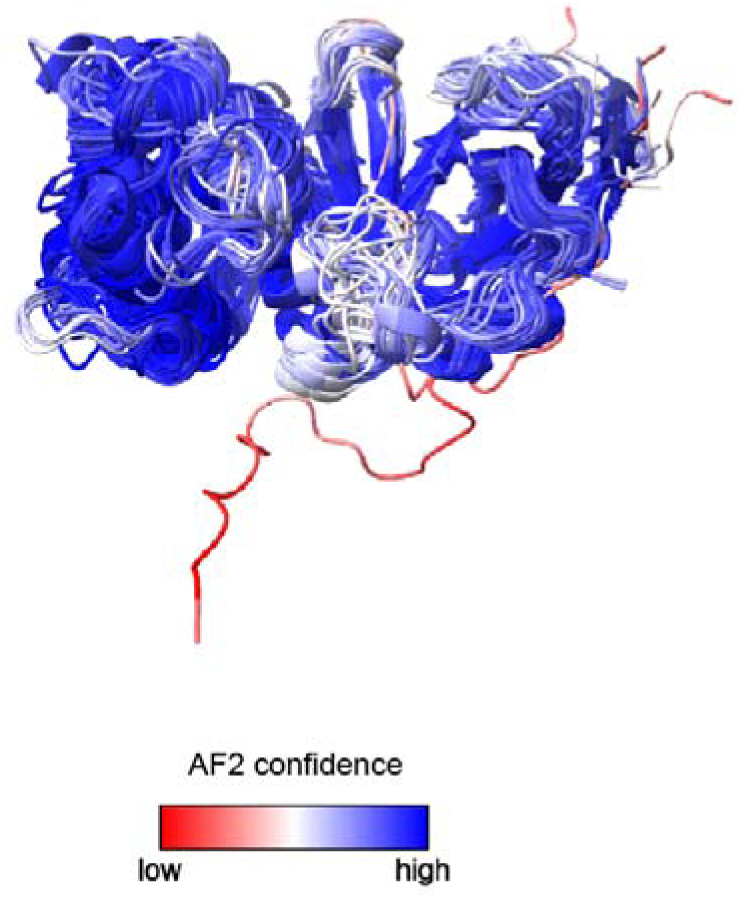
Confidence of Alphafold2 predictions for AcrIF11 homologs.

**Supplementary Figure 4.** Enlarged version of the AcrIF11 phylogeny in Figure 1F. See supplementary file.

**Supplementary Figure 5.**
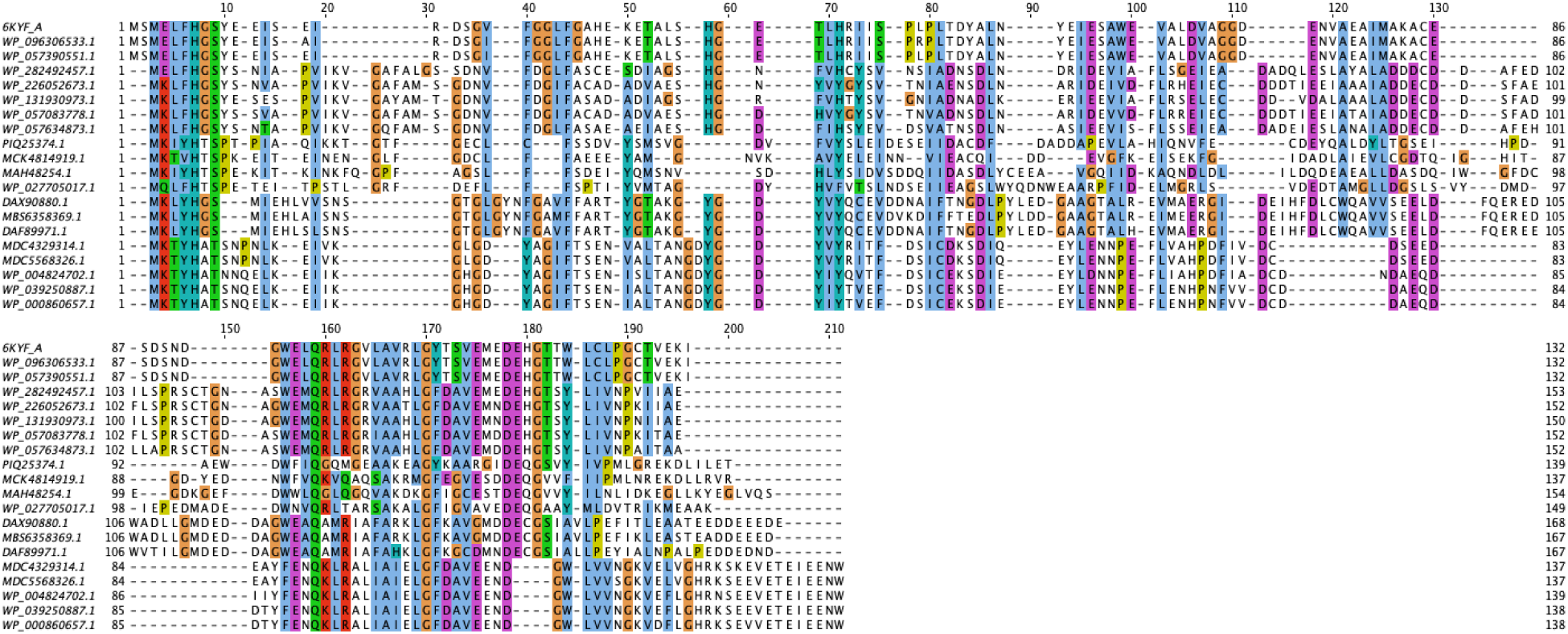
AcrIF11 phylogeny sequence alignment Below is a diverse sampling of the sequence alignment used to build the AcrIF11 phylogeny. Phylogeny construction is discussed in the Methods section.

**Supplementary Figure 6.**
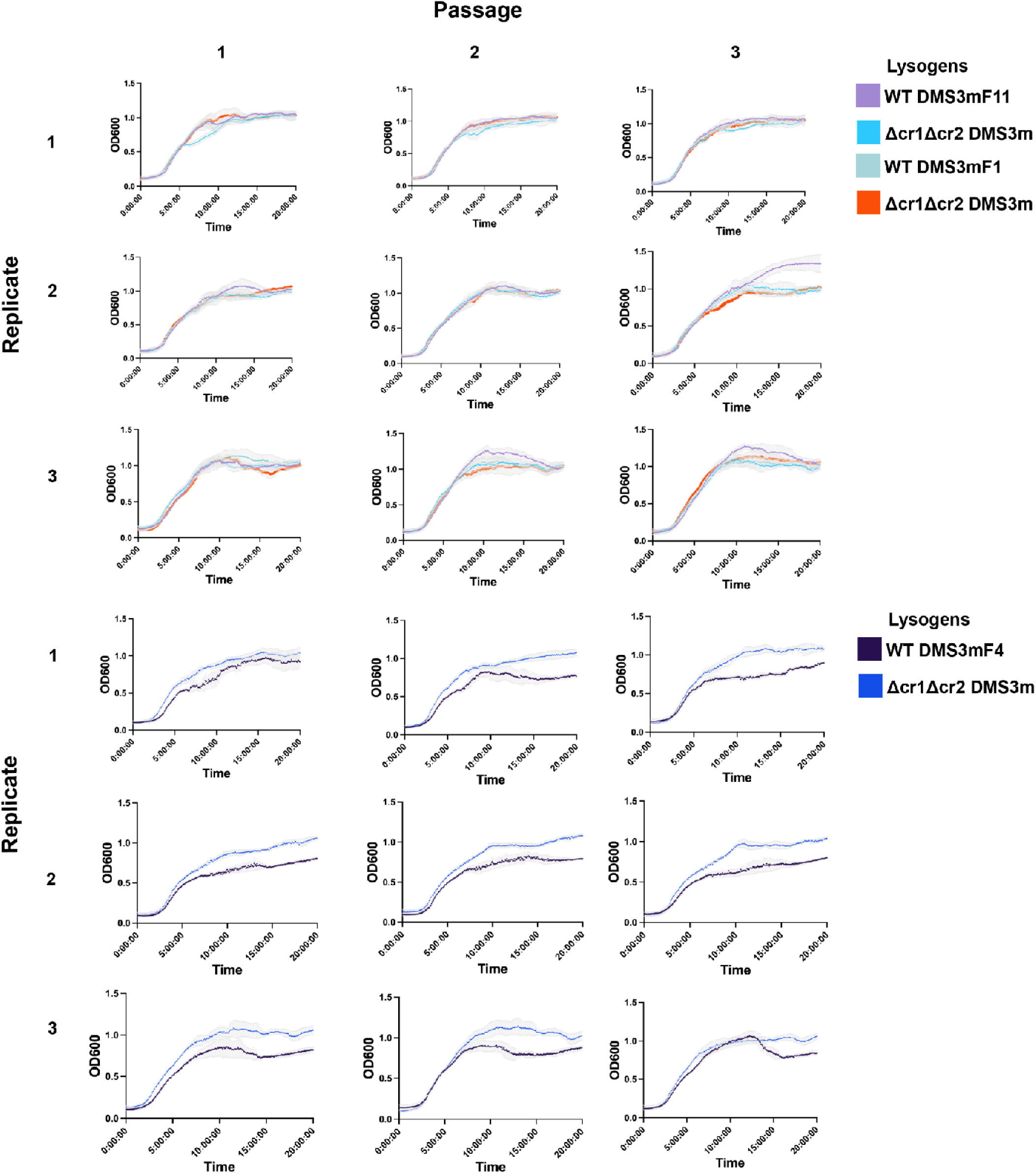
Replicates of lysogen growth experiment

**Supplementary Figure 7.**
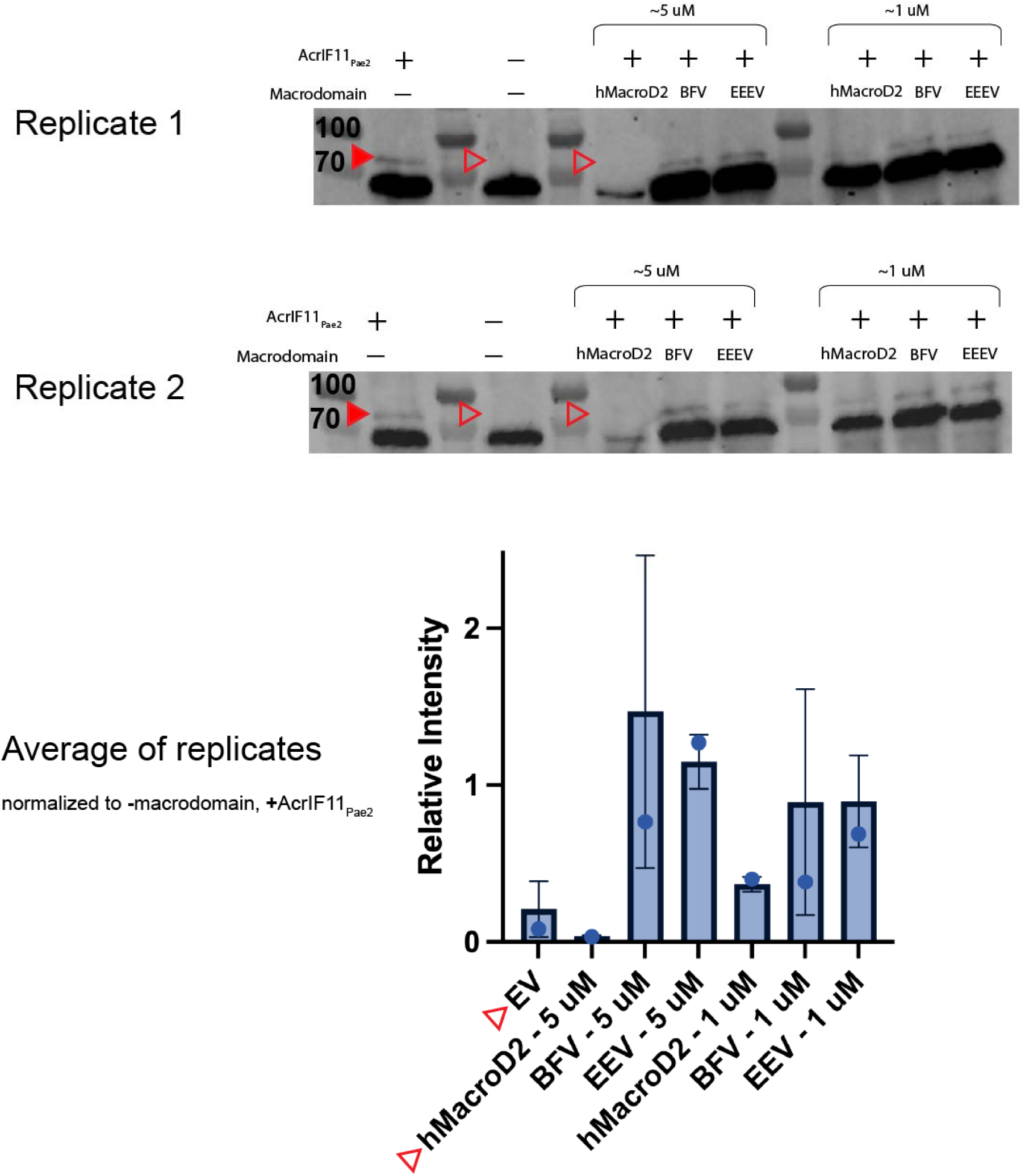
Quantification of macrodomain lysate blot

**Supplementary Figure 8.**
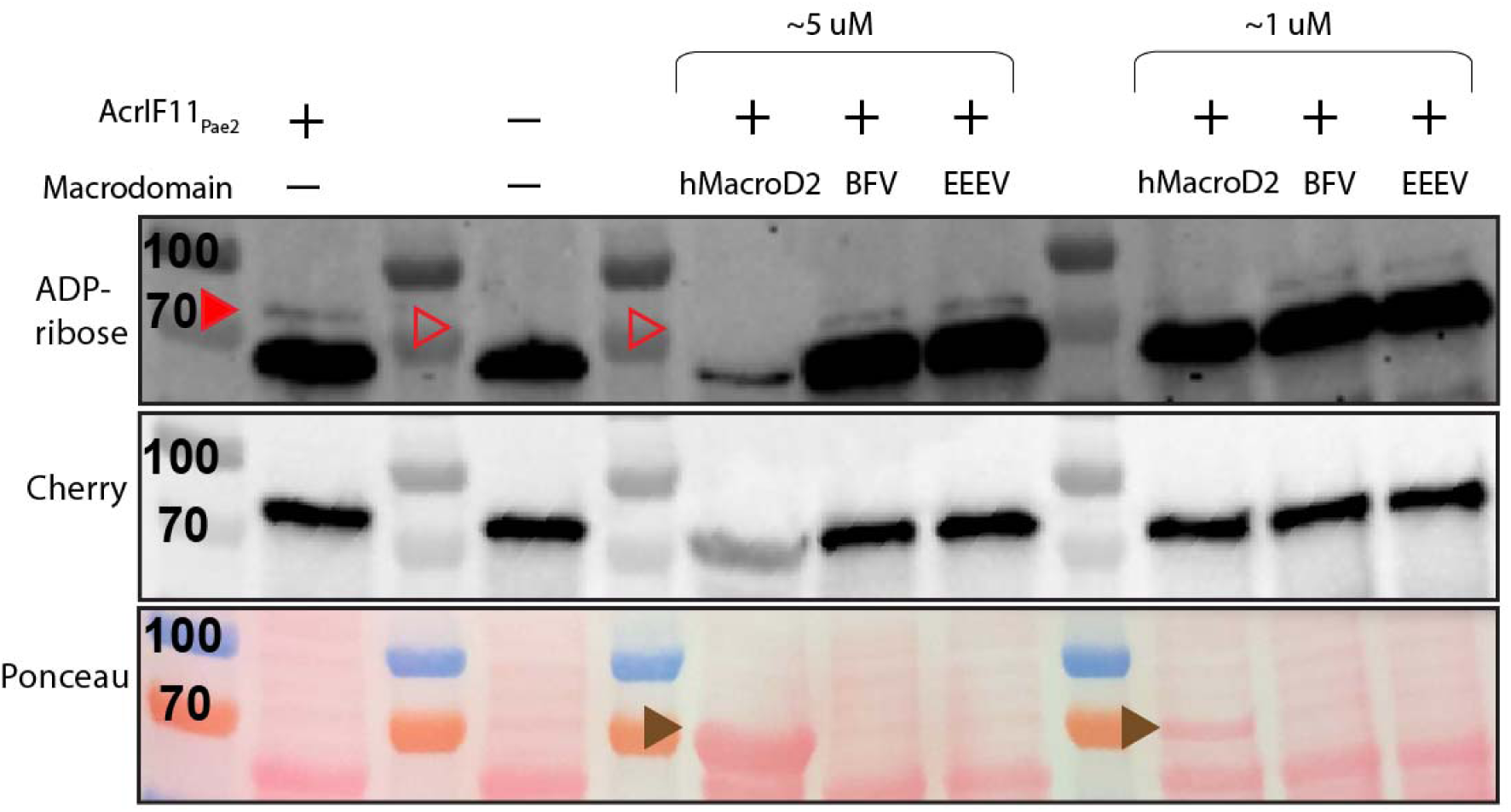
Verification of macrodomain lysate blot loading via Ponceau All labels are the same as in Fig 5B. Brown arrow indicates hMacroD2.

**Supplementary Figure 9.**
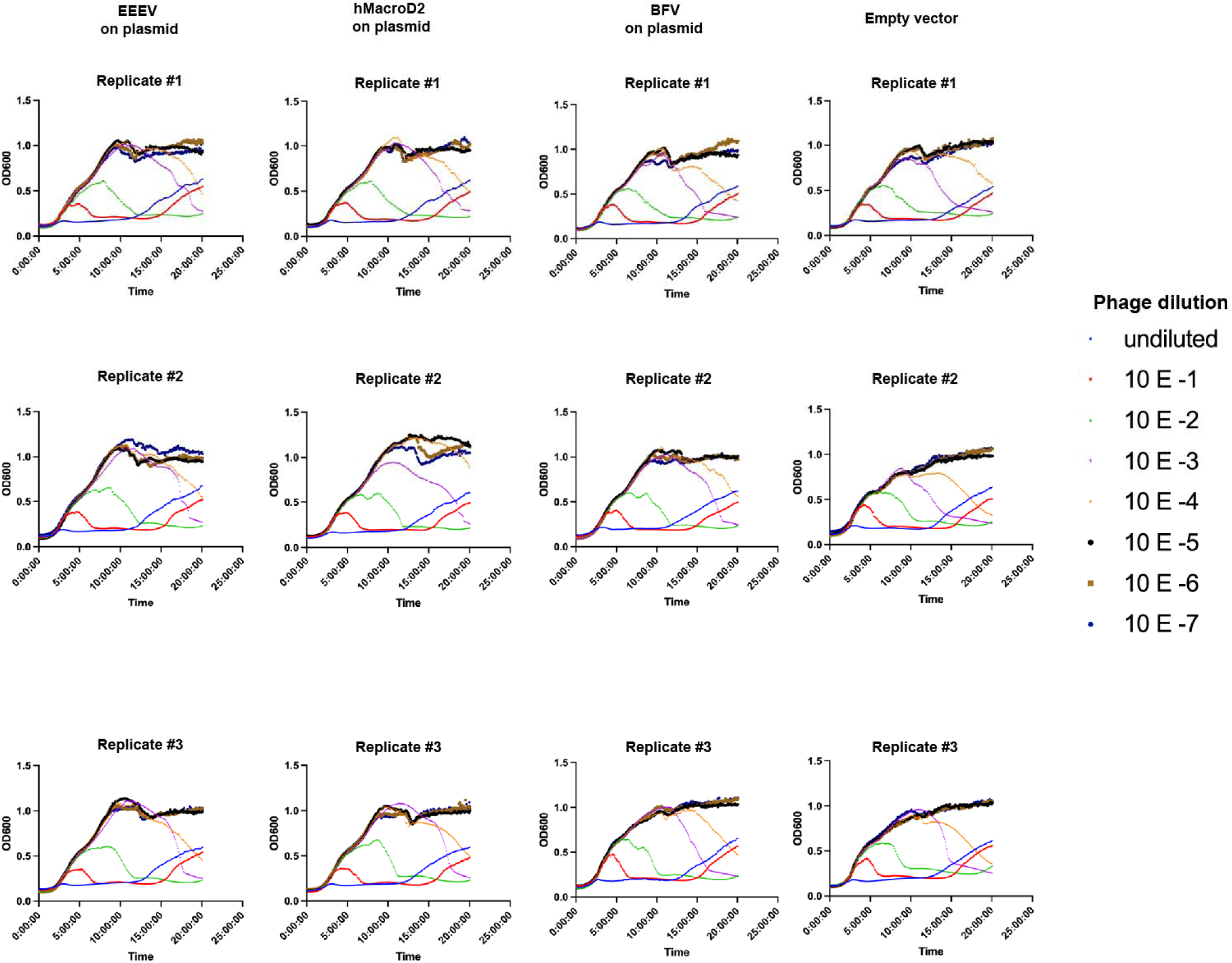
Liquid growth curves of PA14 WT overexpressing non-endogenous macrodomains PA14 WT was infected with DMS3mF11_Pae1_vir, following the lytic infection protocol listed in the Methods section above.

**Supplementary Figure 10.**
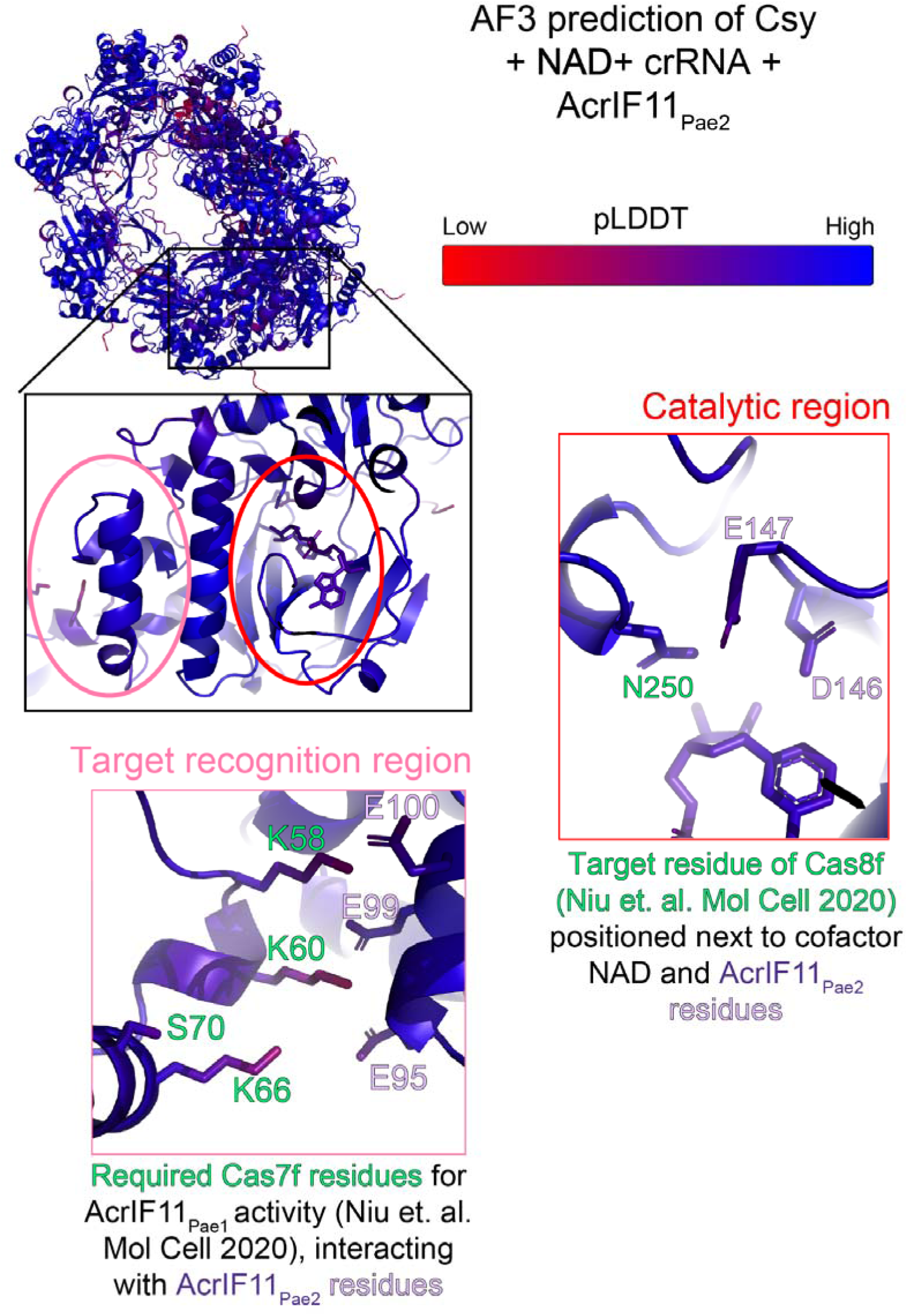
Alphafold3 prediction of Csy complex + NAD + crRNA + AcrIF11_Pae2_ colored by pLDDT

## Author contributions

A.L.B., J.B.D, and J.S.F. conceived the project. A.L.B., J.Y.Z, Y.L. and D.F.C. executed phage & biochemistry experiments. L.T.R. and M.J.S.K. determined the NMR structure. P.B., S.M., M.G.V.S., G.J.C., M.E.D., D.H.S., A.A., R.M.S. provided macrodomains. J.B.D. and J.S.F. supervised experiments and acquired funding. D.F.C., J.B.D, and J.S.F. wrote the manuscript with input from coauthors.

## Acknowledgements

Experiments were funded by awards to J.B.D by the Vallee, Searle, and Kleberg Foundations and to J.S.F. by NIH GM145238 and U19AI171110. D.F.C is supported by the UCSF Discovery Fellowship. D.F.C. and J.S.F would like to thank Dr. Allison Williams for guidance on biochemistry experiments, and Dr. Amy Diallo and Dr. Kliment Verba for BFV and EEEV constructs.

## Competing interest statement

J.B.D. is a scientific advisory board member of SNIPR Biome, Excision Biotherapeutics, and LeapFrog Bio, consults for BiomX, and is a scientific advisory board member and co-founder of Acrigen Biosciences and ePhective Therapeutics. The Bondy-Denomy lab received research support from Felix Biotechnology. A.A. is a co-founder of Tango Therapeutics, Azkarra Therapeutics and Kytarro; a member of the board of Cytomx, Ovibio Corporation, Cambridge Science Corporation; a member of the scientific advisory board of Genentech, GLAdiator, Circle, Bluestar/Clearnote Health, Earli, Ambagon, Phoenix Molecular Designs, Yingli/280Bio, Trial Library, ORIC and HAP10; a consultant for ProLynx, Next RNA and Novartis; receives research support from SPARC; and holds patents on the use of PARP inhibitors held jointly with AstraZeneca from which he has benefited financially (and may do so in the future). J.S.F. is a consultant to, shareholder of, and receives sponsored research support from Relay Therapeutics.

## Notes

https://www.rcsb.org/structure/8DWQ

### Summary of Updates

We now include a new in vivo head-to-head competition assay between AcrIF11 and AcrIF1, improved figure clarity, additional context for macrodomain eraser activity, and expanded phylogenetic and sequence analysis using a broader database. We have also revised the discussion and presentation throughout in response to suggestions on specificity, experimental detail, and formatting.

